# Extrachromosomal DNA as a Causal Instrument for Spatial Multi-Omics

**DOI:** 10.64898/2026.06.02.729483

**Authors:** David W. Craig, Andrei S. Rodin

**Author notes:** Correspondence: David W. Craig.

## Abstract

Spatial transcriptomics, multiplex imaging, and computational pathology now map tissue organization at cellular resolution, but the analyses applied to these data remain correlational. Clustering and co-occurrence statistics describe which features appear together; they cannot say which feature drives the others. We propose a framework for causal inference in spatial multi-omics built on a specific feature of extrachromosomal DNA (ecDNA). ecDNA carries no centromere; it does not attach to the mitotic spindle and partitions randomly to daughter cells at division. Two neighboring cells in the same microenvironment can therefore inherit very different oncogene copy numbers for reasons unrelated to local signaling. Together, these cell-intrinsic randomization properties of ecDNA provide the framework for ecDNA copy number serving as an instrumental variable (IV) separating the effects of oncogene dosage from the downstream cellular effects. In this study, we formalize this within a structural causal model, implement two-stage least squares estimation with sibling-comparison and falsification diagnostics, and provide sensitivity analyses for residual confounding. To benchmark these methods against ground truth, we built CAUSANTA, an agent-based simulator that generates spatial tissue with a known causal graph and stochastic ecDNA inheritance. The framework recovers known oncogene-to-phenotype effects from simulated data, and we outline its application to real tumor sections, including multi-instrument settings where independent ecDNA species carrying different oncogenes enable factorial designs within a single tumor.

## INTRODUCTION

### An Opportunity for Causal Reasoning in Spatial Omics

Spatial transcriptomics and multiplex imaging now resolve tissue states at single-cell resolution, but the analyses applied to them stop at description: clustering, trajectory inference, and correlation-based biomarker discovery.^5^ These methods identify which features co-occur. They do not say which feature is upstream of which, or whether either is upstream of anything at all. The gap matters for translational work: spatial data, on its own, cannot tell us which molecular events drive a tissue state, how that state will evolve, or how it would respond to a targeted intervention ^6-10^.

Current multimodal approaches excel at identifying patterns and correlations but cannot reliably distinguish causes from consequences due to strong influences from the local tissue environment. The core obstacle is confounding by local context. Consider two genetically identical tumor cells sitting a few hundred microns apart in a glioblastoma section. The cell near a perfused vessel proliferates rapidly and is sensitive to cytotoxic therapy. Its neighbor, in a hypoxic pseudopalisade behind a necrotic core, has shifted toward a mesenchymal, quiescent state and tolerates the same drug ^11^. The two cells differ in transcriptional state, morphology, and drug response, yet they are genetically identical: the differences come from oxygen, nutrient gradients, immune contact, and stromal cues ^12^. This is the rule in tissue, not the exception. Methods that pool cells by molecular similarity, including representation-learning approaches developed by us and others ^13,14^, recover spatial motifs but cannot tell whether a transcriptional signature is producing a morphology or being produced by it. Distinguishing these directions is what therapeutic targeting and biomarker development actually require, and current tools do not deliver it.

### Instrumental Variables: From Correlation to Causation

Causal inference provides a formal route from correlation toward causal interpretation^15,16^. More specifically, instrumental variable analysis uses naturally occurring sources of exogenous variation to identify causal effects from observational data under explicit assumptions. In this framing, causal reasoning provides the interpretation layer that converts features into mechanistic hypotheses and counterfactual predictions by explicitly modeling directionality, separating drivers from passengers, and enabling predictions about how biological systems would respond to perturbation rather than simply describing observed states.

Heritable somatic variation propagates across cellular descendants, creating clones with distinct molecular states that share locally comparable microenvironmental contexts, and can serve as a candidate somatic Instrumental <u>V</u>ariable (IV) within a <u>S</u>tructural <u>C</u>ausal <u>M</u>odel (SCM). The logic is that if a somatic event ‘Z’ (the instrument) affects phenotype ‘Y’ only through its effect on gene expression ‘X’, and if Z arose independently of current microenvironmental confounders at the relevant causal point, then the Z→X→Y pathway can isolate causal effects under explicit partly testable assumptions. This approach to causal reasoning, foundational in econometrics and epidemiology, has been underutilized in spatial biology because appropriate biological instruments were not recognized. ^16^ In this work, we identify somatic stochasticity as precisely such an instrument. ^17^

### Somatic Variability Enables Causal Inference from Observational Tissue Data

Somatic variations are post-mitotic changes in DNA sequence that propagate through cell division, generating clonal populations with distinct genetic states. ^**18-20**^ The standard approach for establishing causation relies on controlled intervention, where a variable is perturbed and the resulting effect is measured. In intact tissues, this strategy fails because the required perturbations disrupt the spatial structure under study. Genome editing, drug treatment, and genetic knockouts alter tissue organization, while dissociation removes spatial context entirely. Organoid systems lack key features of native microenvironments, and xenograft models introduce confounding from species differences. ^17,21^ Alternative causal approaches face similar limitations: Mendelian randomization depends on germline variants shared by all cells in a tissue, providing no leverage for distinguishing causal relationships among neighboring cells.

Concurrent independent work has introduced a “Somatic-IV” framework that uses patient-level somatic mutations and copy number alterations as instruments for survival outcomes to identify candidate cancer drivers^22^. We introduce a framework that is complementary in operating at single-cell resolution within one tumor, deriving instrument validity from a biophysical mechanism (acentric ecDNA segregation) rather than statistical analogy to Mendelian randomization, and estimating continuous causal effect sizes for known structural edges rather than identifying drivers from cohort survival.

Our central supposition is that tissues themselves contain a solution for causal Inference. Somatic processes generate heritable variation among nearby cells through stochastic events that occur largely independently of local context. X-chromosome inactivation patterns, mitochondrial DNA mutations, and somatic mosaicism all introduce variation in gene dosage or allelic state among neighboring cells (Table 1). This variation propagates across cellular descendants, creating clones with distinct molecular states sharing locally comparable microenvironmental contexts. This variation acts as a natural source of quasi-random perturbation, enabling instrumental variable analysis while preserving native tissue architecture. ^23^

**Table 1.**
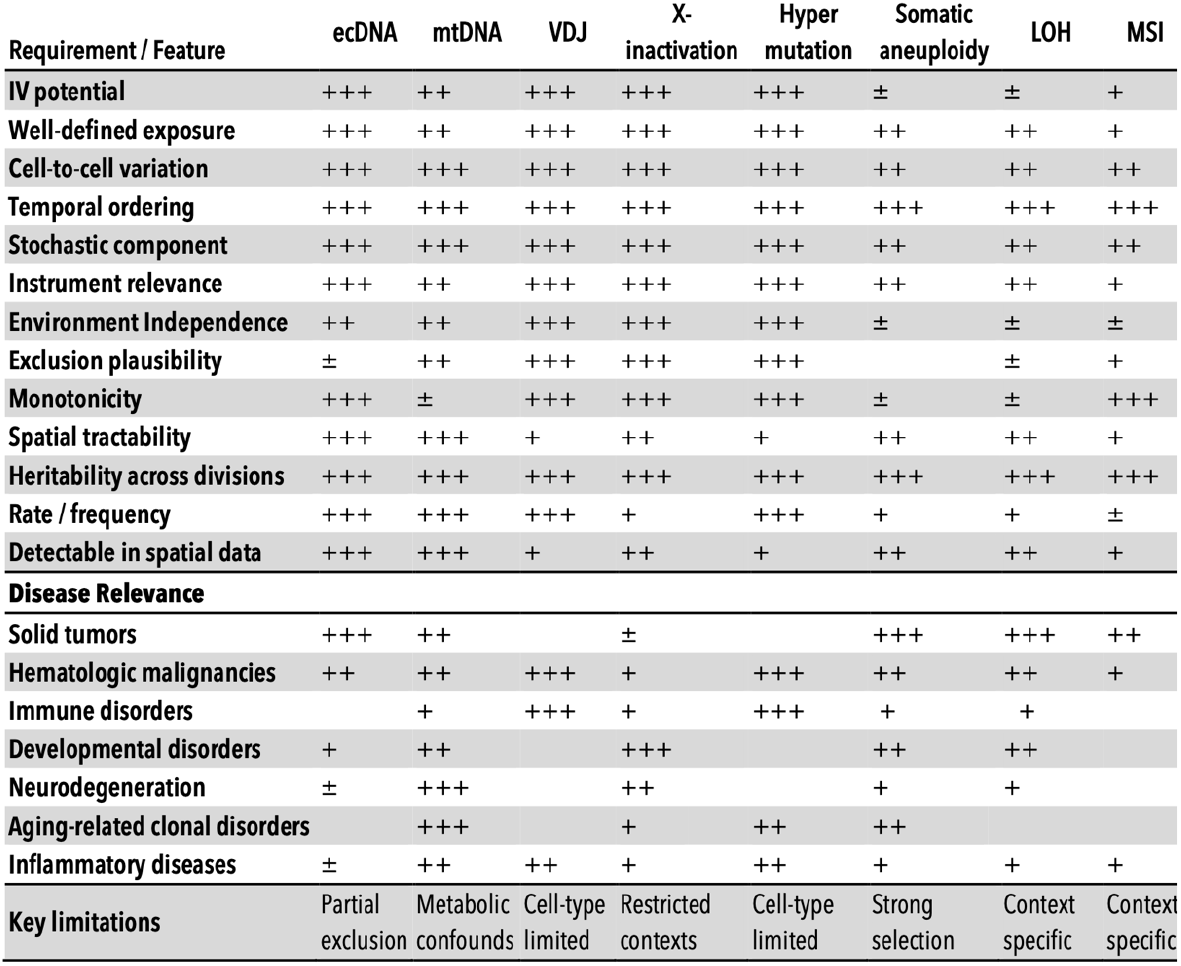
Comparative evaluation of somatic biological variations as instrumental variables for causal inference in disease contexts. This framework qualitatively summarizes various somatic features according to criteria central to instrumental variable analyses, including definability of exposure, stochasticity of cell-to-cell variation, temporal ordering, heritability, spatial detectability, and adherence to core instrumental variable assumptions (relevance, independence from confounding, and exclusion) ^1-3^. Practical considerations such as event frequency, known limitations, instrumental variable utility, and disease specificity are also indicated. Qualitative ratings reflect relative suitability: (+++) strong, (++) moderate, (+) limited, (±) context-dependent, and (–) generally unsuitable.

### Extrachromosomal DNA as a Proof-of-Principle System for Spatial Causal Inference

Extrachromosomal DNA (ecDNA) is the cleanest known instance of somatic variation with the properties the <u>S</u>omatic <u>I</u>nstrumental <u>V</u>ariable (SIV) framework requires. ecDNAs are circular DNA molecules, typically megabase-scale, that carry oncogenes and regulatory elements but lack a centromere ^24-27^. Without a centromere they do not attach to the mitotic spindle, and at division they partition to daughter cells in a process well approximated by binomial sampling (Figure 1). Single-cell sequencing, FISH tracking, and live-cell imaging all support this stochastic inheritance ^28^. The consequence at tissue scale is that two neighboring cells in the same microenvironment can inherit different ecDNA copy numbers, and therefore different oncogene dosages, purely because of how the molecules partitioned at the last mitosis. This is the natural experiment the framework needs. ecDNAs were first described as “double minutes” in the 1960s and characterized cytogenetically by Mark and colleagues in 1970 ^29^. Pan-cancer analyses now identify ecDNA in roughly 15–17% of tumors, with substantially higher frequencies in aggressive subtypes including glioblastoma ^24^. ecDNA drives elevated oncogene transcription through accessible chromatin architecture and accelerates adaptation to selective pressure, including therapy ^26,30^. Our group reported the first long-insert whole-genome reconstruction of an ecDNA element in 2013 ^4^ and recently mapped the spatial architecture of co-occurring *EGFR* and MDM2 ecDNAs producing distinct tumor compartments in glioblastoma ^31^.

**Figure 1.**
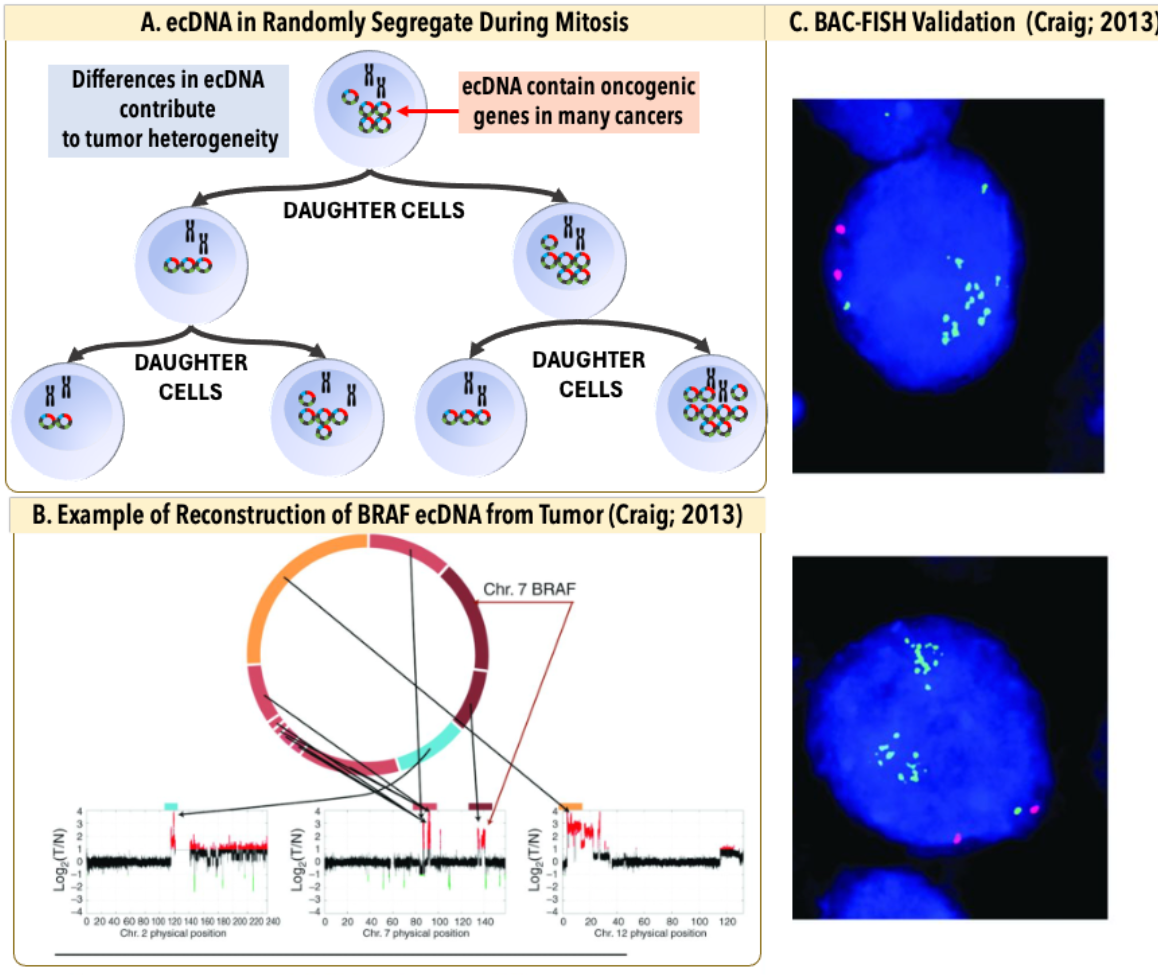
Extrachromosomal DNA as an instrumental variable for causal inference in cancer. **(A)** ecDNAs are circular DNA elements located outside chromosomes in tumor nuclei that frequently carry oncogenes and regulatory sequences. Their random segregation during mitosis generates stochastic copy number variation that is independent of local microenvironmental states, while increased copy number drives elevated transcription, satisfying key requirements for an instrumental variable. **(B)** Example reconstruction of ecDNA architecture published by Craig et al., 2013 ^4^ using long-insert mate-pair sequencing, showing CN at breakpoints linking segments from chromosomes 1, 7, and 12. Arrows indicate amplified junctions. (C) BAC-FISH validation of BRAF-containing ecDNA, with locus-specific BAC probes labeled in green and chromosome 7 centromeric probes labeled in red.

Three formal conditions are required for valid IV inference, and ecDNA can be assessed against each ^15^. **Relevance**: the instrument must move the exposure substantially. ecDNA segregation generates copy-number variation spanning roughly 1 to over 100 copies per cell, and two decades of work, most recently from the Mischel laboratory, establishes that oncogene copy number on ecDNA tracks closely with expression and downstream pathogenicity ^24,30^. **Independence**: the instrument must arise independently of unmeasured confounders of the exposure-outcome relationship. Mitotic segregation is a physical sampling event that occurs before the daughter cells re-establish transcriptional, metabolic, and signaling states; recent live-cell and lineage-tracing work supports independence at the segregation event ^28^. **Exclusion**: the instrument must affect the outcome only through the exposure. This is the strongest and most biologically vulnerable assumption, because ecDNA can carry passenger regulatory elements, alter nuclear organization through hub formation, and modify mitotic dynamics in ways that are not captured by oncogene dosage alone. We do not assume exclusion holds. We probe it: the framework includes falsification analyses using negative-control exposures, sensitivity analyses bounding the effect of plausible violations, and triangulation against sibling-comparison estimators that rely on different assumptions.

### ecDNA *EGFR*-Encoded Glioblastoma as a Demonstration Model

Glioblastoma is the demonstration system. IDH-wildtype GBM has a high frequency of ecDNA amplification and frequently co-amplifies *EGFR* and MDM2 on what appear to be largely independent ecDNA species. The resulting spatial pattern has been characterized at single-cell resolution: proliferative transcriptional programs concentrate in *EGFR*-rich regions, and hypoxia-response programs concentrate in MDM2-rich regions ^31^. The biology is consistent with two opposite causal stories. In one, *EGFR* amplification drives proliferation through ERBB signaling, and proliferating cells happen to be perivascular for unrelated reasons. In the other, perivascular cells proliferate because they have oxygen and nutrients, and ecDNA copy number tracks proliferation through positive selection rather than driving it. Standard correlative analyses cannot distinguish these, and the distinction is not cosmetic: an *EGFR*-directed therapy will reduce tumor growth only in the first scenario. The same logic applies to MDM2-driven p53 inhibition in the hypoxic niche, where causal direction determines whether MDM2 inhibitors meaningfully alter the niche population. The IV framework can address both.

### Framework and Scope

This paper does four things. First, we formalize the Somatic Instrumental Variable (SIV) framework as a Structural Causal Model (SCM) for spatial tumor biology, with explicit estimators, falsification diagnostics, and sensitivity analyses. Second, we introduce CAUSANTA (Causal Analysis Using Somatic And Neighborhood Tissue Architecture), an agent-based spatial simulator that generates virtual tumor sections with a known causal graph and stochastic ecDNA inheritance, providing ground truth against which IV and causal-discovery methods can be benchmarked. Third, we show that the framework recovers known oncogene-to-phenotype effects in simulation and lay out the design for applying it to real tumor sections. Fourth, we extend the approach to tumors carrying multiple ecDNA species with different oncogene payloads, where approximately independent segregation enables factorial designs within a single section ^31^.

## RESULTS

### Simulating Tumor Sections with Known Causal Structure

We built the CAUSANTA-TME (TME, tumor microenvironment) simulation engine to answer a specific causal question: can stochastic ecDNA inheritance separate oncogene-driven cellular phenotypes from the spatial microenvironment that usually confounds them? The simulator (shown in Figure 2) was designed as a ground-truth benchmark, not as a predictive model of glioblastoma. In each run, the causal graph, ecDNA inheritance rule, hypoxia-*EGFR* coupling, and *EGFR*-to-phenotype effect sizes were fixed before analysis. This allowed every estimator to be scored against a known answer. These ground-truth sections are the substrate for the analyses that follow, and as shown in examples in Figure 3.

**Figure 2.**
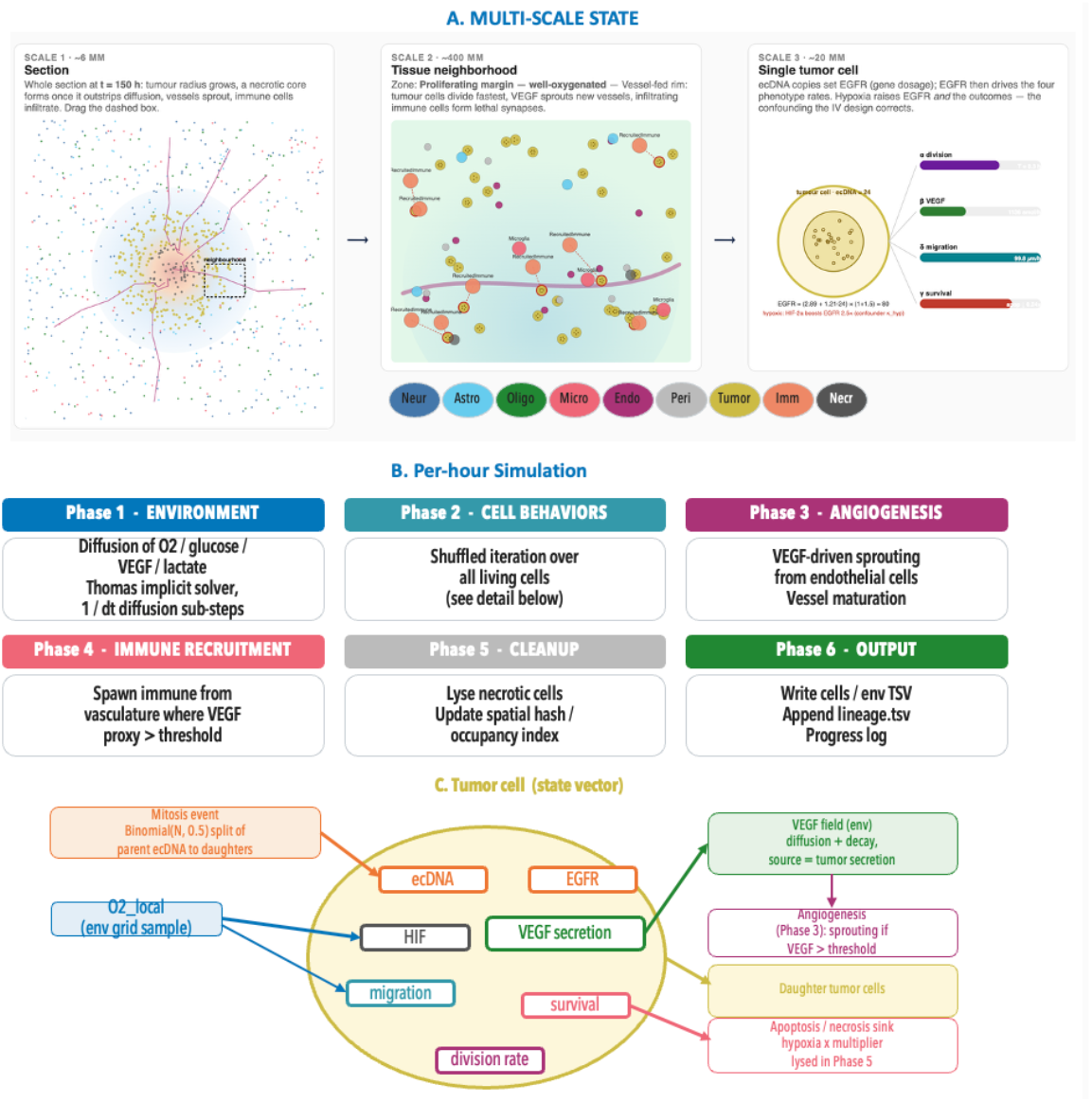
Architecture of the CAUSANTA-TME simulator. **(A)** CAUSANTA-TME simulates a coupled multi-scale tumor system, beginning with a tissue-scale domain and extending to individual cells. The model includes subclonal structure, tumor and non-tumor cell types, vascular regions, immune context, and normal tissue compartments. Each simulated cell is assigned morphology, local microenvironment, *EGFR* expression, hypoxia status, parentage, and ecDNA copy number. **(B)** These cell-level features are organized into spatial neighborhoods and converted into experimental measurement layers, including H&E morphology, FISH, Visium-like spots, and single-cell readouts. **(C)** The simulator retains the underlying ground truth for each feature, including subclone identity, ecDNA-driven *EGFR* expression, hypoxia, proliferation, apoptosis, and necrosis. These linked outputs allow benchmarking of whether an analysis can recover causal structure rather than simply detecting spatial correlation. In the benchmark workflow, simulated ecDNA, *EGFR* expression, morphology, and spatial context are used as inputs to the PC causal discovery algorithm, with graph structure recovery performance evaluated against the known ground truth.

**Figure 3.**
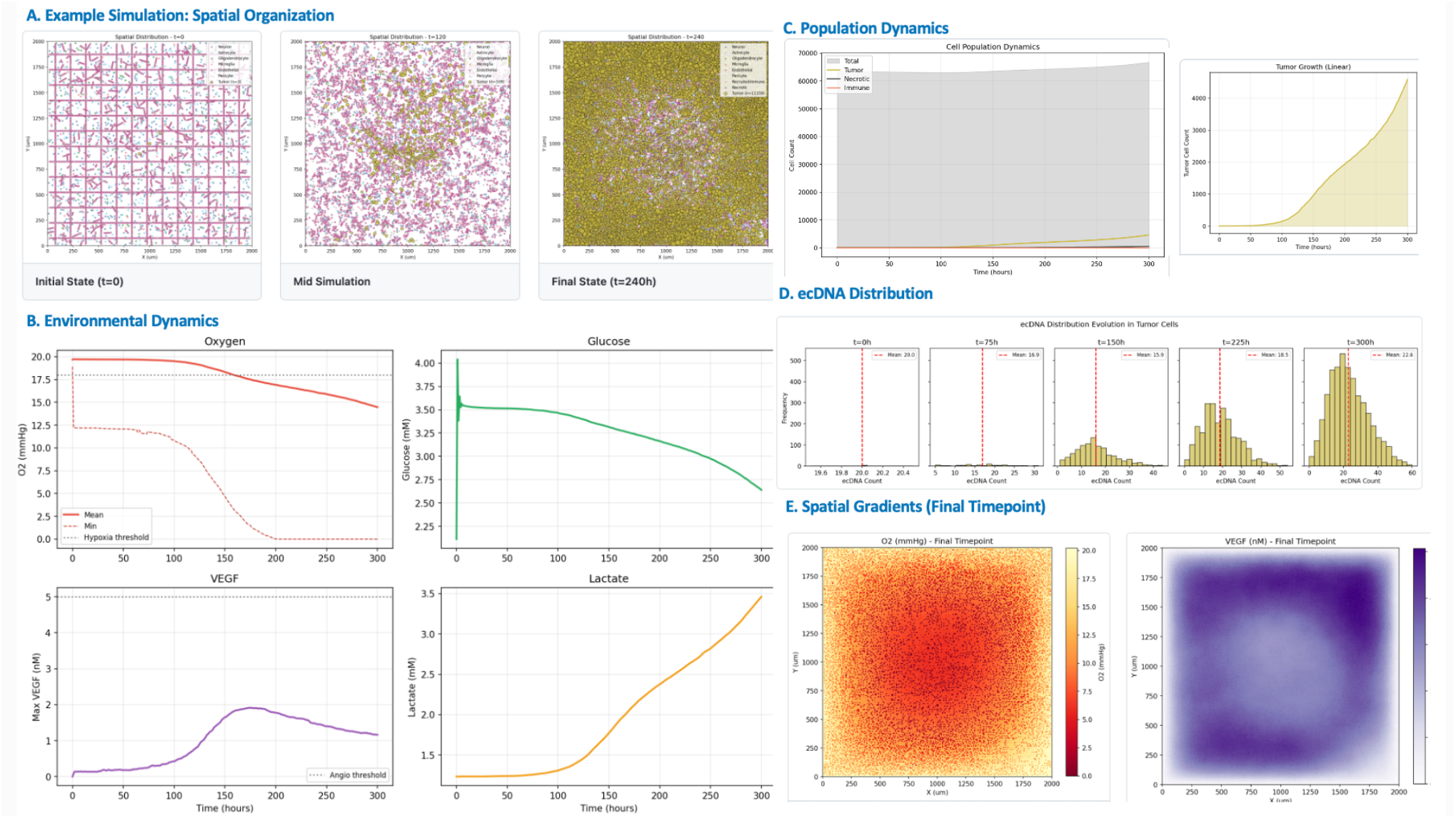
Tissue dynamics. **(A)** Spatial organization of the simulated tumor at three timepoints — initial (t = 0), mid, and final (t = 300 h) — showing outward radial growth from the seeded founders into the surrounding normal-tissue compartment. **(B)** Environmental dynamics over the time course: local oxygen and glucose decline toward the growing tumor core while *VEGF* and lactate accumulate, driven by hypoxic *VEGF* secretion and Warburg-type lactate production. **(C)** Tumor population growth on linear and log scales, showing the exponential-then-saturating trajectory. **(D)** Distribution of ecDNA copy number per tumor cell at successive timepoints, which broadens through stochastic mitotic segregation to span roughly 0–60 copies. The simulator retains the ground truth for every feature, enabling benchmarking of whether an analysis recovers causal structure rather than mere spatial correlation. **(E)** Steady-state spatial gradients at the final timepoint (oxygen and *VEGF* concentration fields).

The benchmark was organized around three expectations. First, if ecDNA segregation behaves as a somatic instrument, daughter-cell copy-number allocation should be random with respect to local oxygen, glucose, position, lineage age, and other microenvironmental variables. Second, ecDNA copy number should strongly predict *EGFR* expression, creating a high-powered first stage. Third, when hypoxia influences both *EGFR* and downstream phenotypes, ordinary least squares should be biased, while two-stage least squares and sibling comparisons should recover the planted causal effects. When the hypoxia-to-*EGFR* edge is removed, the OLS-IV gap should collapse.

We tested these predictions in a 2 x 3 x 5 design: two spatial scales (2 mm and 6 mm), three hypoxia-*EGFR* coupling strengths (baseline, reduced, and removed), and five random seeds per scale-coupling cell, for 30 simulations. Each run generated a spatial tumor section with cells, vasculature, oxygen, glucose, lactate, *VEGF*, ecDNA copy number, *EGFR* expression, hypoxia status, and behavioral phenotypes. The analyses below follow the same diagnostic ladder needed for real tissue: biological validation, instrument relevance, instrument independence, effect recovery, falsification, robustness, and spatial effect mapping.

### CAUSANTA generated the expected tumor ecology and ecDNA heterogeneity

The simulated tumors reproduced the features that make spatial tumor data difficult to interpret. Tumors expanded radially from seeded founders, consumed oxygen and glucose, accumulated *VEGF* and lactate, developed hypoxic and necrotic regions, and generated angiogenic feedback. These behaviors satisfied the pre-specified validation targets and established that the benchmark contained a spatially structured microenvironment rather than independent cell-level observations.

The same runs produced the molecular heterogeneity required for the SIV design. Starting from founder cells carrying 20 *EGFR*-ecDNA copies, stochastic replication and mitotic segregation broadened the copy-number distribution as tumors grew. In the representative 6 mm baseline run, tumor-cell ecDNA copy number ranged from 0 to 64 copies, with a median of 22, a mean of 24.0, an interquartile range of 15 to 31, and a 5th to 95th percentile range of 7 to 46 (Figure 3). Thus, cells in the same simulated tumor carried a broad gradient of oncogene dosage.

This heterogeneity is the first biological reason ecDNA can be useful for causal inference. The analysis does not require comparing different patients or different tumors. It requires neighboring or closely related cells inside one tumor that differ in the instrument value. CAUSANTA generated exactly that structure: shared tissue context with stochastic cell-to-cell differences in ecDNA copy number.

### Stochastic ecDNA segregation produced a natural experimen

The first formal requirement was independence at the segregation event. CAUSANTA records every mitosis, including the parent copy number, the replicated ecDNA pool, and the number of copies assigned to each daughter. This made it possible to test the randomization mechanism directly.

Across 23,205 division events in the canonical 6 mm baseline run, daughter ecDNA allocation matched the Binomial(N, 0.5) expectation. The mean daughter fraction of the replicated pool was 0.4994, statistically indistinguishable from 0.5 (one-sample t test, t = -1.01, p = 0.31), and the observed variance was consistent with the per-cell binomial expectation. Across the additional 6 mm baseline seeds, more than 100,000 division events showed the same pattern. S-phase replication also matched the configured rule: the mean post-S/pre-S copy-number ratio was 1.95, corresponding to a per-copy replication probability of 0.948 against the configured value of 0.95.

The daughter fraction was not associated with candidate confounders measured at division. In the canonical 6 mm baseline lineage, daughter fraction showed no meaningful association with x-coordinate (r = -0.006, p = 0.34), y-coordinate (r = 0.002, p = 0.75), generation number (r = 0.003, p = 0.70), parent ecDNA count (r = 0.002, p = 0.76), simulation time (r = 0.003, p = 0.60), distance from tumor center (r = 0.005, p = 0.43), or parent-cell oxygen at division (r = -0.0005, p = 0.94).

These were the expected negative results. The tumor itself was highly spatially structured, but the act of ecDNA allocation at mitosis was not spatially biased. This distinction is essential: oxygen, hypoxia, and regional tumor architecture can remain confounders of *EGFR*-expression analyses while ecDNA copy number remains a valid instrument for the randomized component of *EGFR* dosage.

### ecDNA copy number was a strong first-stage predictor of *EGFR*

The second requirement was relevance. ecDNA copy number strongly predicted *EGFR* expression across the factorial benchmark. Fitting the structural first-stage model *EGFR* = X_base + κZ + κ_hyp M(X_base + κZ) + ε recovered κ = 1.22 ± 0.004 across the five 6 mm baseline seeds, matching the configured per-copy *EGFR* contribution of κ = 1.21.

The hypoxia-dependent term was recovered as well. In the baseline condition, hypoxic cells followed an *EGFR*-copy-number slope of approximately 3.03, matching the theoretical value 1.21 x (1 + 1.5) = 3.025. In the removed-confounder condition, where κ_hyp = 0, the simple *EGFR* ∼ Z regression recovered an intercept of 2.91 ± 0.17 and a slope of 1.21 ± 0.01, within sampling error of the configured values. Across all 30 simulations, first-stage F-statistics ranged from 29,057 to 149,960, far above the conventional weak-instrument threshold of 10.

This result rules out a weak-instrument explanation for the analyses that follow. ecDNA copy number moved *EGFR* expression strongly and predictably. Differences between OLS and IV therefore reflect the modeled confounding structure, not lack of instrument power.

### The confounding problem was visible before causal estimation

The simulated sections reproduced the central ambiguity of spatial tumor biology. A high-*EGFR* cell with high migration, high *VEGF*, or altered survival could be responding to ecDNA dosage, responding to a hypoxic niche, or both. In the baseline scenario, hypoxia increased *EGFR* through the HIF-mediated coupling and also altered downstream phenotypes through proliferation arrest, migration programs, and *VEGF* induction. A regression of phenotype on *EGFR* therefore mixed the dosage effect with the niche effect.

The factorial design provided an internal falsification test for this problem. We expected OLS and IV to diverge when hypoxia acted as a shared cause of exposure and outcome. We expected them to converge when that shared-cause pathway was removed. This prediction was borne out across scales and seeds.

### Causal discovery recovers the projected ground-truth graph in a curated benchmark

Effect estimation assumed the causal graph; we next asked whether the graph itself can be recovered from the same observational data without pre-specifying which edges to look for. We applied the PC constraint-based causal discovery algorithm with significance level α = 0.05 and maximum conditioning-set size k = 3 to ecDNA copy number, *EGFR* expression, hypoxia, oxygen-linked variables, proliferation, migration, *VEGF*, and survival without specifying a single target edge. The prediction is that PC should recover the ecDNA → *EGFR* → phenotype skeleton with correct identifiable edge orientation, should exclude direct ecDNA → phenotype edges (the exclusion restriction the simulator enforces), and should not generate spurious edges at the chosen significance level (Figure 4).

**Figure 4.**
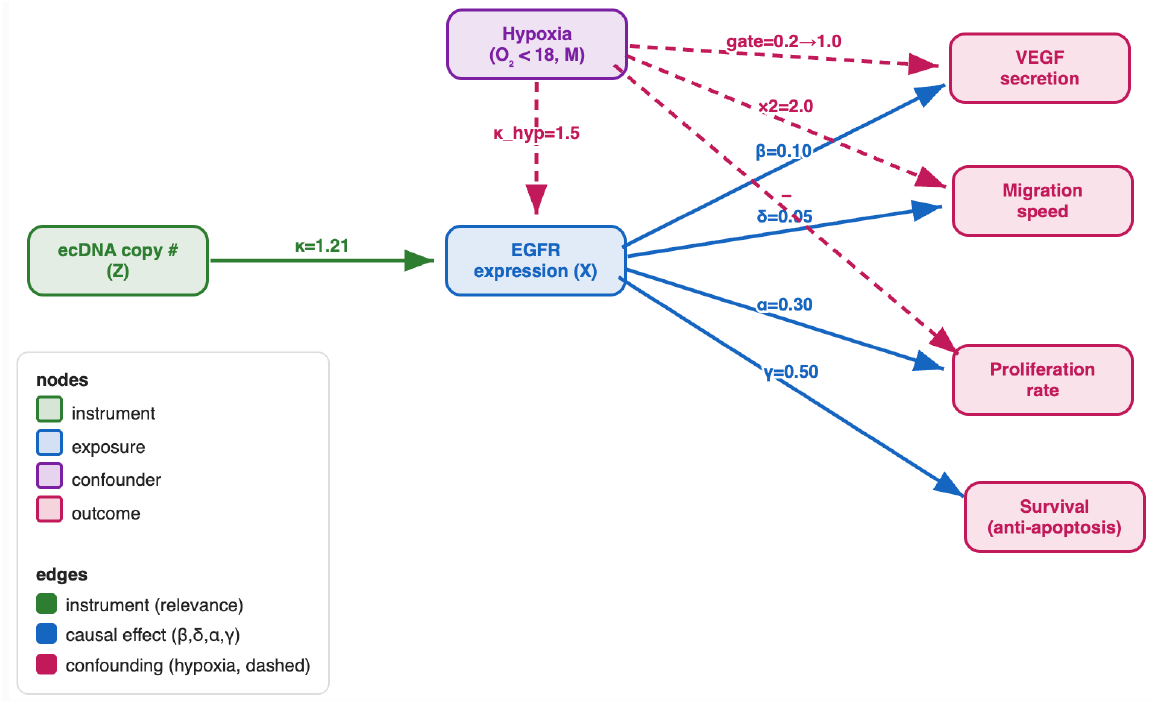
Directed acyclic graph encoding the somatic instrumental variable framework as implemented in CAUSANTA. ecDNA is a somatic instrumental variable: random mitotic segregation (Z) drives *EGFR* (X); hypoxia (M) confounds X and the outcomes. 2SLS using Z recovers the true effects; OLS is biased.

At 2 mm baseline, PC recovered seven edges with correct orientation: ecDNA → *EGFR, EGFR* → Proliferation, *EGFR* → Migration, *EGFR* → *VEGF*, O2 → Proliferation, O2 → Migration, and Hypoxia → *VEGF*. Two edges were not fully recovered. The *EGFR* → Survival edge appeared in the skeleton, but its orientation was unresolved within its Markov equivalence class. The O2 → Hypoxia edge collapsed during preprocessing, where the two variables were merged into a single node. No spurious edges were inferred at α = 0.05. Importantly, PC did not infer direct ecDNA → phenotype edges, consistent with the exclusion restriction encoded in the simulator. Structural Hamming Distance was 0; precision, recall, and F1 were 1.00 against the projected ground-truth graph used for scoring Across the full factorial design (n = 5 seeds × 6 cells), F1 ranged from 0.790 to 1.000, with the highest performance at 2 mm baseline (1.000 ± 0.000), where the collider structure is most informative. Recall was perfect except at 6 mm reduced (0.880); precision varied more widely (0.663 at 2 mm removed to 1.000 at 2 mm baseline). The wider seed-to-seed spread in the reduced and removed scenarios reflects the known difficulty of resolving collider structure when an incoming edge is weak or absent.

### IV recovered the planted causal effects, whereas OLS tracked the hypoxic niche

Using ecDNA copy number as the instrument for *EGFR* expression, 2SLS recovered the configured *EGFR* effects on downstream phenotypes. Structural inversion recovered the *VEGF* effect β = 0.100 and the migration effect δ = 0.050 across the multi-seed benchmark, matching the planted values. This is the primary positive-control result: when the ground truth is known, the IV estimator returns it.

Naive regression did not. In the 6 mm baseline condition, linear OLS estimated the *EGFR*-to-*VEGF* effect as 4.11 ± 0.13, while 2SLS estimated 3.64 ± 0.11, an OLS inflation of +12.5%. In the 2 mm baseline condition, OLS exceeded IV by approximately +8.7% (Figure 5). The larger bias in the larger tumor matched the expected biology: broader spatial context created stronger hypoxic structure, and hypoxia was a shared cause of *EGFR* elevation and *VEGF* induction.

**Figure 5.**
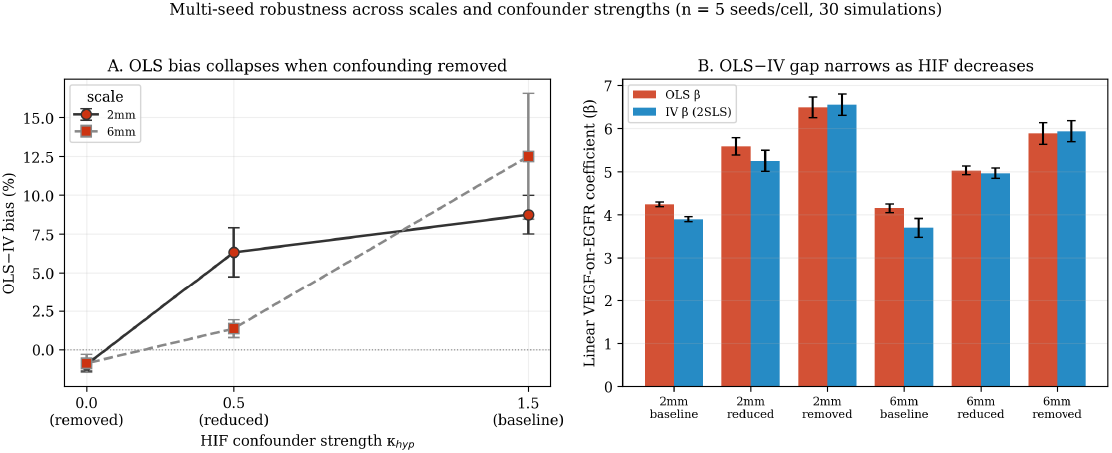
Multi-seed robustness analysis (n = 5 per cell, 30 simulations total). **(A)** OLS-vs-IV bias percentage as a function of HIF confounder strength, with error bars showing seed-to-seed standard deviation. Bias collapses to within one percent of zero (−1.0% ± 0.4 at 2mm; −0.9% ± 0.6 at 6mm) when the confounding edge is removed (HIF = 0.0), at both scales tested. The tight reliability bounds rule out seed-specific accidents. **(B)** OLS β (red) and IV β (blue) per cell with error bars from seed variance. The gap between OLS and IV narrows monotonically as HIF decreases; at HIF = 0.0 the two estimators agree to within 1% at both scales. IV and OLS show comparable seed-to-seed variability across cells (e.g. equal SD of 0.056 at the 2mm baseline).

The OLS-IV gap changed exactly as predicted when the confounding pathway was manipulated. Under baseline hypoxia-*EGFR* coupling, OLS exceeded IV by +8.7% at 2 mm and +12.5% at 6 mm. Under reduced coupling, the bias dropped to +6.3% at 2 mm and +1.4% at 6 mm. Under removed coupling, OLS became statistically indistinguishable from IV, with bias near zero at both scales (-1.0% ± 0.4 at 2 mm and -0.9% ± 0.6 at 6 mm; Figure 6).

**Figure 6.**
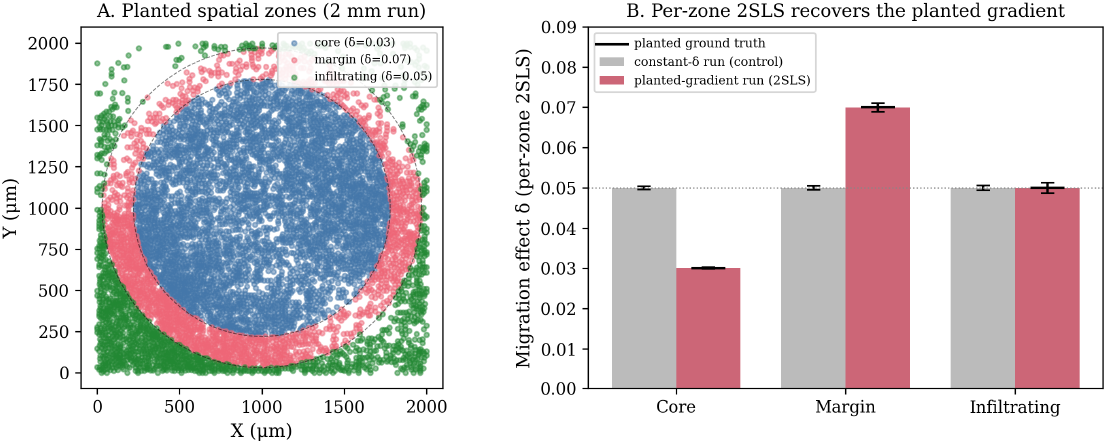
Spatial heterogeneity of causal effects. **(A)** Planted spatial zones (2 mm gradient run). Tumor cells are colored by the concentric zone they occupy, defined by distance from the tumor-seed centroid: core (< 780 µm; planted δ = 0.03), margin (780–970 µm; planted δ = 0.07), and infiltrating (> 970 µm; planted δ = 0.05); dashed circles mark the zone boundaries. **(B)** Per-zone 2SLS estimate of the *EGFR*-on-migration effect δ for two runs. In the constant-δ control (grey), the per-zone estimate is flat at δ = 0.050 in every zone, so the framework does not fabricate a gradient where none exists. In the planted-gradient run (red), the identical per-zone 2SLS recovers δ = 0.030 / 0.070 / 0.050 for core / margin / infiltrating, each matching the planted ground truth (black markers) to the third decimal. The estimator is therefore both specific (flat when the effect is uniform) and sensitive (it recovers true effect modification when present).

This is the most important benchmark result. The bias was not introduced by the IV estimator. It appeared when the structural causal model said OLS should be confounded, weakened when the confounding edge was weakened, and disappeared when the edge was removed. ecDNA-IV therefore estimated the *EGFR*-driven component of the phenotype, while OLS estimated a mixture of *EGFR* dosage and spatial niche.

### Sibling comparisons independently validated the estimates

We next tested the same causal effects using sibling comparisons. At a mitotic event, two daughter cells share a parent, spatial origin, and immediate history, but they can inherit different ecDNA copy numbers through binomial allocation. Comparing siblings removes every confounder they share, including unmeasured confounders.

The sibling design agreed with the IV estimates. For migration, the sibling/2SLS estimate was δ = 0.050 across the 30 simulations, matching the planted value. For *VEGF* secretion, the corresponding estimate was β = 0.100, also matching the planted value. Agreement among the planted ground truth, population IV, and sibling contrasts provides stronger validation than any estimator alone.

### Falsification and robustness analyses supported the IV assumptions

We challenged the estimates using negative controls and hidden-confounding sensitivity analyses. Rosenbaum bounds were off-scale over the tested range: for both the migration and *VEGF* structural effects, no hidden binary confounder with odds-ratio distortion up to Γ = 50 made the IV estimates lose significance at α = 0.05.

E-values provided a second scale for the same question. At a 10-copy ecDNA contrast, the E-value was 3.07 for migration and 1.34 for *VEGF*, with essentially identical confidence-interval E-values. The *VEGF* E-value is smaller because of the effect scale, so the more direct falsification test is whether ecDNA copy number spuriously tracks spatial position.

It did not. ecDNA copy number did not predict cell x-coordinate (β = 0.002, p = 0.81), y-coordinate (β = -0.001, p = 0.89), or distance from tumor center (β = 0.008, p = 0.34). These null placebo outcomes show that the instrument was not simply a surrogate for where a cell sat in the simulated tumor.

### Spatial and temporal benchmarks showed where the framework is informative

A spatial causal estimator should not invent spatial effect modification when none exists, and it should recover spatial effect modification when it is planted. We tested both using the *EGFR*-to-migration effect δ estimated separately in tumor core, margin, and infiltrating zones (Figure 6).

In the constant-effect negative control, per-zone 2SLS returned δ = 0.050 in all three zones, with confidence intervals within ±0.0004 of the planted value. In the planted-gradient run, where δ was set to 0.03 in the core, 0.07 at the margin, and 0.05 among infiltrating cells, the same estimator recovered δ = 0.0302, 0.0699, and 0.0500, respectively. The method was therefore specific when the effect was uniform and sensitive when the effect was heterogeneous.

We also re-ran IV estimation and PC causal discovery at intermediate timepoints to ask whether causal recovery depended on tumor age (Figure 7). Within the simulated window, PC causal discovery improved as tumors grew. At 6 mm baseline, F1 rose from 0.738 at t = 100 h to 0.954 at t = 300 h. At 2 mm baseline, F1 rose from 0.707 at t = 120 h to 1.000 at t = 240 h. Increasing the sample size more than compensated for decreasing variance in the binary hypoxia indicator.

**Figure 7.**
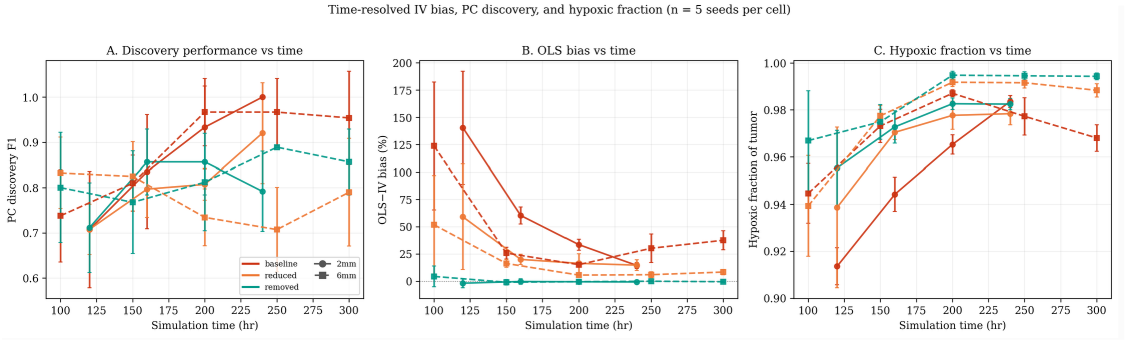
Time-resolved IV and PC causal discovery performance (n = 5 seeds per cell). **(A)** PC causal discovery F1 vs simulation time, with error bars showing seed-to-seed SD. F1 generally increases as the tumor population grows and the structural edges become more reliably detectable (non-monotone in the reduced/removed cells, where the collider edge is weak or absent). **(B)** OLS-vs-IV bias percentage vs time. Bias is largest in the smallest early tumors and falls as the tumor matures; the 6 mm baseline trajectory is non-monotone, reaching a minimum near t = 200 hr before rising again. **(C)** Hypoxic fraction (fraction of tumor cells with O_2_ below the 18 mmHg HIF threshold) vs time. Hypoxic fraction is already > 85% at the earliest measured timepoint and never drops below this level — but this is by construction of the model, not a methodological artifact: tumor cells consume O_2_ heavily, and the simulator only initializes tumor seeds in the parenchymal interior, so the moment the seed cells divide, they create a local hypoxic micro-environment. The full simulation domain remains a mix of normoxic (vascular regions and tissue periphery) and hypoxic (tumor core) tissue throughout. The relevant point is that even when all tumor cells are hypoxic, the PC algorithm can still recover the (is_hypoxic → *EGFR*_expression) edge because (i) the binary indicator still has > 0 variance in the tumor population (264 of 8,458 cells are normoxic at t = 300 hr 6 mm baseline; even one normoxic cell with low *EGFR* is informative against the dosage-only prediction), and (ii) IV does not require the confounder to be detectable at all. Solid lines = 2 mm runs (circles); dashed lines = 6 mm runs (squares). Colors: red = baseline (HIF = 1.5), orange = reduced (HIF = 0.5), green = removed (HIF = 0).

OLS bias was largest early, when spatial gradients were forming and sample sizes were smaller. At 2 mm baseline, OLS exceeded 2SLS by +140.5% at t = 120 h but by +14.5% at t = 240 h. At 6 mm baseline, the bias fell from +123.8% at t = 100 h to approximately +15% near t = 200 h, then rose to +37.5% at t = 300 h. Bias and recoverability were therefore dynamic properties of the tumor state, not fixed properties of the estimator.

### Summary of the Results

Across the benchmark, the observed results matched the expected signatures of a valid SIV framework. ecDNA segregation was random at mitosis, ecDNA copy number strongly predicted *EGFR*, OLS was biased when hypoxia confounded exposure and outcome, the OLS-IV gap collapsed when that confounding edge was removed, and both 2SLS and sibling comparisons recovered the configured effects.

The framework also passed the checks that make the result interpretable rather than merely positive. Placebo outcomes were null, sensitivity analyses did not overturn the estimates, causal discovery recovered the intended graph with high F1, and per-zone estimation distinguished true spatial effect modification from a flat causal map. These results establish CAUSANTA as a ground-truth benchmark for spatial causal inference and show that ecDNA copy number can function as a somatic instrument under the assumptions encoded in the simulator.

## DISCUSSION

We have shown that the stochastic segregation of ecDNA at mitosis can be exploited as a Somatic Instrumental Variable (SIV) for causal inference in spatial tumor biology, and we have built CAUSANTA, an agent-based simulator that embeds a known causal graph into realistic tumor tissue to benchmark the framework against ground truth. In simulation, the SIV approach recovered the configured causal parameters within sampling error while ordinary least squares overestimated them by 8.7 to 140.5%. A factorial sensitivity analysis across confounder strengths and tumor scales showed that the OLS bias is proportional to the strength of the Hypoxia → *EGFR* confounding pathway and collapses to within ∼1% of the IV estimate when that pathway is removed, providing direct empirical confirmation that the bias is the bias the structural causal model predicts, not an artifact of the estimator. The remainder of this section discusses what this enables, what it does not yet establish, and what real-tumor validation will require.

### Biological and Translational Implications

Driver-versus-passenger discrimination is one of the oldest open problems in cancer genomics. The standard formulation, an alteration is a driver if it contributes causally to malignancy, presupposes a way of measuring causal contribution that the field has rarely been able to deliver. SIV provides one such way at the level of individual ecDNA-encoded oncogenes and individual phenotypes. In our simulations, the configured causal effect of *EGFR* amplification on migration (δ = 0.05) is less than half the naive OLS estimate (β_OLS = 0.12). A correlation-based prediction of how much migration would fall under *EGFR* inhibition would therefore overstate the achievable benefit by more than two-fold. The same logic applies to any therapeutic decision based on cross-sectional correlation between an amplified oncogene and a downstream phenotype.

Spatially resolved causal estimates can in principle generate therapeutically actionable maps, but only where genuine effect modification exists. In the current simulator the migration coefficient is spatially constant, so the recovered effect map is flat (a negative control confirming the method does not fabricate a gradient). In real glioblastoma, where effect modification across niches is biologically plausible, the same per-zone 2SLS would resolve true spatial gradients while mitigating bias from the spatial structure of confounders. The mechanistic candidates that could drive a real core-versus-margin difference are contact inhibition restraining the *EGFR*-driven motility program in the dense core, and reduced ECM resistance combined with stronger selective pressure for migration at the margin, both testable in slice-culture and intravital-imaging models. The proliferation effect, by contrast, peaks in the perivascular core where oxygen and nutrients are available, and antiproliferative agents should concentrate there. The general claim is that ecDNA-driven oncogene effects are spatially structured, and effective therapy may need to be spatially structured to match.

### Multi-Instrument Extensions: Factorial Designs within a Single Tumor

Many tumors carry more than one ecDNA species. Glioblastomas with co-amplified *EGFR* and MDM2 on apparently independent ecDNA molecules are well documented ^31^, and combinations involving PDGFRA, CDK4, and MYC are also common in GBM. To the extent that two ecDNA species segregate independently of each other, their copy numbers in any given cell are approximately uncorrelated, and the resulting joint distribution mimics a 2 × 2 factorial design embedded in a single tumor section. Multi-instrument IV analysis can then estimate the causal effect of each oncogene while using the other as a check on the exclusion restriction: if both instruments yield consistent causal estimates for a shared downstream outcome, exclusion is additionally supported for that outcome; if estimates diverge, at least one instrument violates exclusion, and the analysis can identify which one. A tumor of ∼10,000 cells with moderate ecDNA prevalence will populate each factorial cell with hundreds of cells, sufficient for stable estimation. Whether real co-occurring ecDNA species are independent enough for this design is itself an empirical question; if their inheritance is correlated through shared origin (chromothripsis) or hub-mediated co-segregation, the design becomes a fractional factorial with partial aliasing, recoverable but with reduced power.

### Generalization: Other Somatic Instrumental Variables

The SIV concept extends to any heritable somatic variation that arises by a mechanism independent of the local microenvironment. Several candidates are tabulated in Table 1; three deserve mention here.

Mitochondrial heteroplasmy is the closest analogue to ecDNA. mtDNA molecules partition stochastically across daughter cells at mitosis, and cells carrying a heteroplasmic pathogenic variant inherit different mutation loads across sibling lineages ^32^. The relevant instrument is the cell-level heteroplasmy fraction; the exposure is the corresponding change in mitochondrial respiration, ROS generation, or mtDNA-encoded protein synthesis; downstream phenotypes include metabolism and apoptotic threshold. Two complications distinguish mtDNA from ecDNA: the segregation process is not strictly binomial because mitochondria fuse and fragment between divisions, and mtDNA mutation rates can themselves be elevated by microenvironmental stress (hypoxia, ROS), so the at-the-segregation-event independence condition needs case-by-case empirical support.

Chromosomal instability in aneuploid tumors creates ongoing missegregation of whole chromosomes or chromosome arms. The instrument is per-cell chromosome copy number; the exposure is per-arm gene dosage; downstream phenotypes are arm-level fitness effects. The signal is noisier than ecDNA segregation because missegregation events are rarer and the resulting copy-number variance is smaller, but the same machinery applies.

Copy-neutral loss of heterozygosity (CN-LOH) arises by mitotic recombination or non-disjunction-plus-reduplication. It is binary, irreversible, and propagates clonally, providing a binary instrument for allele-specific expression and haploinsufficiency. The timing of the LOH event also encodes lineage information that aids causal discovery, particularly when paired with other clonal markers.

Across these examples, the operational question is the same one we asked for ecDNA: does the candidate variation move the exposure (relevance), arise independently of the relevant confounders at the moment of the event (independence), and affect the outcome only through the named exposure (exclusion)? Stochastic DNA methylation at metastable epialleles can be framed in the same way, but the independence-from-environment condition is the weakest of the candidates here because methylation rates are themselves environmentally sensitive; we mention it as the boundary case rather than a recommendation.

### Relation to Prior Work

The IV framework originated in econometrics to address confounding in observational policy analysis ^33^. The canonical instruments are external shocks (lottery outcomes, the Vietnam draft, weather, geographic distance to a service) that influence the exposure of interest without otherwise affecting the outcome. Our contribution is to identify a biological process, ecDNA segregation at mitosis, that meets the same logical specification but at the cellular rather than the individual level.

Mendelian randomization transposed the IV framework to genetic epidemiology by using germline variants as instruments for modifiable exposures, transforming the field’s ability to make causal claims from cohort data ^21^. Germline variants take a single value across every cell of an individual, however, which makes them useless for within-tumor or within-tissue causal questions. SIV is the natural extension of MR to the somatic and intra-tissue setting: where MR exploits randomization at conception, SIV exploits randomization at mitosis. The consequence is that one tumor section contains thousands of independent realizations of the randomization, where one MR study contains one realization per participant.

Recent causal-inference work in spatial and single-cell biology has taken three forms: perturbation-based validation of correlational spatial associations (Perturb-seq and its spatial extensions); causal-discovery algorithms applied to spatial graphs to recover putative regulatory edges; and concurrent independent work introducing a patient-level Somatic-IV framework ^22^ that uses cohort-level somatic mutations and copy number alterations as instruments for survival outcomes. Our contribution is distinct on three axes: (i) the instrument is biophysical (acentric ecDNA segregation), not statistical analogy to MR; (ii) the unit of randomization is the cell, not the patient; and (iii) the estimands are continuous causal effects on specified spatial phenotypes, not driver identification from cohort survival.

### Limitations and Assumptions

The exclusion restriction is the strongest assumption the framework places on the biology. It requires that ecDNA affect downstream phenotypes only through the expression of the genes they carry, and there are at least four candidate mechanisms by which it could fail. ecDNA hubs cluster in the nucleus and can engage in trans-regulatory interactions that activate loci on other ecDNA molecules or on chromosomes ^28^, producing pleiotropic effects that bypass the named exposure. ecDNAs can sequester transcription factors, chromatin modifiers, or replication machinery in their hub microenvironment. The metabolic and replicative cost of carrying tens to hundreds of extra copies of an oncogene-containing molecule is non-zero and is not captured by gene-expression mediators. Cytoplasmic ecDNA fragments can trigger cGAS-STING innate immune sensing ^26^, potentially affecting tumor-immune interactions independent of the cargo gene. The current evidence suggests these effects are small at physiological copy numbers, but they have not been systematically excluded. The strongest empirical test we know of is to compare cells with matched cargo-gene expression but different ecDNA copy numbers, which is most cleanly achieved by comparing ecDNA-amplified cells to homogeneously staining region (HSR)-amplified cells of the same gene, since HSR amplifications produce equivalent gene dosage but lack the hub-formation, segregation, and immune-sensing properties of ecDNA.

The independence assumption requires that the segregation event itself is independent of unmeasured confounders. Our simulator enforces this by construction; in real tumors it is an empirical claim. Three plausible violations exist. Asymmetric cell division, which is documented in stem-cell biology and may occur in tumor cells under specific niche signals, could bias ecDNA allocation toward the daughter inheriting a particular polarity axis. Differential survival of ecDNA-high versus ecDNA-low daughters across microenvironments creates a post-segregation selection that imprints a correlation between copy number and niche; the segregation event itself remains independent, but the observed population fails the conditional-independence test if cells are sampled long after the event. Cosegregation of ecDNAs with non-ecDNA cellular asymmetries (old-versus-new centrosome inheritance, midbody-tagged components) would couple ecDNA inheritance to other lineage features. The key empirical diagnostic is whether the daughter ecDNA fraction correlates with measured microenvironmental variables at the moment of division. This requires live-cell imaging of dividing cells with ecDNA reporters, or lineage tracing that captures sister cells immediately post-division, neither of which is yet standard in clinical tumor samples.

Applying SIV to real data requires ecDNA copy number at single-cell resolution (via FISH or amplicon sequencing), gene expression co-measured in the same cells (via spatial transcriptomics), phenotypic outcomes measurable from spatial data (morphology, neighbors, protein markers), and lineage information for sibling comparisons (via clonal barcoding or live imaging). Current platforms provide the first three measurements but rarely the fourth. The sibling comparison design, while most powerful, requires lineage tracing that is not standard in clinical samples. The population IV design works without lineage information but has weaker identification.

CAUSANTA is a benchmark, not a predictive model of glioblastoma. It operates in 2D, uses phenomenological reaction-diffusion physics on a coarsened grid, simulates a single ecDNA species, assumes fixed causal-edge weights, and excludes therapy-induced clonal selection and somatic evolution within a section. These are deliberate simplifications appropriate to the benchmarking purpose, but they mean CAUSANTA cannot be used to predict patient-level outcomes, drug responses, or tumor trajectories. What it establishes is that the SIV framework recovers configured causal effects when the underlying generative process is the one the framework assumes. Whether the framework recovers real causal effects in real ecDNA-positive tumors is an empirical question that real-tumor data must answer.

Similarly, high graph-recovery results should be interpreted as biologically curated benchmarking against known ground truth; unconstrained causal discovery from raw spatial omics will require more general causal discovery methods that handle latent confounding, spatial dependence, measurement noise, and mixed data types.

### Future Directions

Three lines of work follow directly from this paper:

The first is real-tumor validation. *EGFR*-amplified glioblastoma is the natural starting point: ecDNA prevalence is high, the *EGFR* pathway is therapeutically actionable, and the spatial architecture of ecDNA distribution has been characterized at single-cell resolution ^31^. A validation study requires fresh-frozen tumor sections with co-registered ecDNA copy number (DNA-FISH or amplicon sequencing) and spatial transcriptomics, ideally paired with orthogonal perturbation data such as *EGFR*-inhibitor response in matched models. The first deliverable is a comparison of the 2SLS estimate of the *EGFR*-on-migration effect in real tumors against (a) the OLS estimate from the same data and (b) the perturbation-derived effect from matched models.

The second is multi-instrument exclusion testing. Tumors carrying *EGFR*-ecDNA and a second co-occurring ecDNA species (MDM2, MYC, CDK4, PDGFRA) allow the same downstream phenotype to be estimated using two instruments. Concordant estimates support exclusion for both. Discordant estimates localize the violation. The *EGFR*/MDM2 co-amplification described by Webb and colleagues ^31^ is a candidate substrate; the empirical question of whether the two species segregate independently in those tumors is itself the first test.

The third is the sibling-comparison design, which is the most powerful form of SIV analysis because it controls every confounder that sibling cells share at the moment of division. Implementing this requires lineage tracing of dividing cells with ecDNA-resolved readouts. CRISPR-tagged ecDNA reporters combined with live-cell or longitudinal organoid imaging are the obvious approach. We are not aware of an existing pipeline that combines ecDNA copy-number tracking with sibling-cell phenotyping in either organoid or tissue contexts; building one is a non-trivial technical commitment but yields the cleanest data the SIV framework can act on.

## Conclusion

The stochastic segregation of ecDNA at mitosis turns one of the most aggressive features of cancer biology into a tool for causal inference. By treating ecDNA copy number as a Somatic Instrumental Variable within a structural causal model, we have a principled way to separate the effects of an amplified oncogene from the effects of where the cell carrying it happens to sit. In simulation, this framework recovers configured causal effects within sampling error while naive regression overestimates them by 8% to 140%. Whether it does the same in real tumors is the next question, and the answer is testable now. Moving spatial biology from correlation to causation does not require new technology; it requires applying the right inferential framework to the data the field is already generating.

## METHODS

### Simulation Engine

CAUSANTA-TME is an agent-based simulator coupled to a reaction-diffusion solver for diffusible substrates. Its purpose in this paper is to generate spatial tumor sections with a fully specified causal graph and configured causal-edge weights, so that any IV or causal-discovery method applied to the resulting data can be scored against known ground truth. The simulator is not intended as a predictive model of glioblastoma; it is a benchmarking substrate. All numerical defaults reported in this section are those used for the multi-scale runs reported in Results.

### Spatial Domain and Temporal Discretization

Simulations run on a rectangular 2D domain at configurable size. Three scales were used: 1000 × 1000 µm, 2000 × 2000 µm, and 6000 × 6000 µm (1 mm, 2 mm, 6 mm). Cell positions are stored at single-micron resolution. Diffusible substrates are solved on a coarser regular grid with spacing h_env = 10 µm (e.g., a 600 × 600 grid for the 6 mm domain). The main simulation time step is Δt = 1 hr; diffusion is sub-stepped at Δt_diff = 0.01 hr (36 s) for numerical stability. Total simulation duration ranged from 168 hr (1 mm runs) to 300 hr (6 mm runs).

### Environment Fields and Reaction-Diffusion

Each environment field *E*_*k*_(***x***, *t*) ∈ {O_2_, glucose, VEGF, lactate} evolves according to a reaction-diffusion PDE:

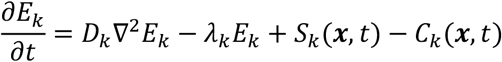

where *D*_*k*_ is the substrate’s effective diffusion coefficient, *λ*_*k*_ is its first-order decay rate, *S*_*k*_ is the source term from vasculature and cellular secretion, and *C*_*k*_ is the sink from cellular consumption. Substrate parameters are summarized in Table 2.

**Table 2.**
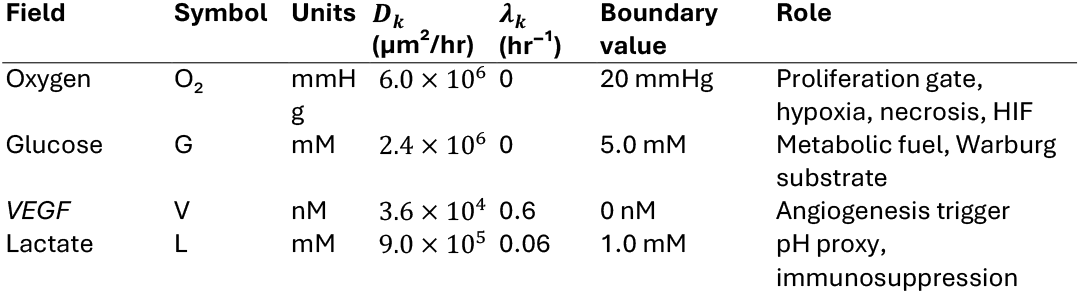
Environment substrates and their PDE parameters. Diffusion coefficients are converted to μm^2^/hr from physiological μm^2^/min reference values. Boundary values for the multi-scale runs correspond to the modestly hypoxic tissue regime used in the 6mm publication configuration.

#### Source and sink terms

Oxygen and glucose are supplied at vascular positions in proportion to local vascular density:

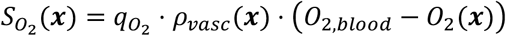

with transfer coefficient 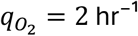 hr^−1^ and arterial reference *O*_2,*blood*_ = 40 mmHg. Cellular consumption acts as a point sink at the voxel containing each cell:

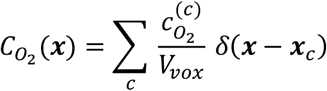

with V_vox = h^2^_env. Per-cell oxygen consumption rates are 30,000 amol/hr for neurons and 72,000 amol/hr for tumor cells. *VEGF* is secreted only by hypoxic cells (O_2_(x_c) < θ_hyp, where θ_hyp = 18 mmHg). Lactate is produced by tumor cells in proportion to glucose consumption (Warburg metabolism).

Dirichlet boundary conditions fix each field to its tissue-baseline value (Table 2), representing the surrounding tissue as an infinite reservoir. Diffusion is solved by implicit Locally One-Dimensional (LOD) operator splitting: for each substrate at each diffusion sub-step we solve an implicit tridiagonal system in x via the Thomas algorithm (one solve per row), then in y (one solve per column), apply source/sink and decay as a separate algebraic step, enforce boundary conditions, and clamp values to physical ranges. Implicit LOD is unconditionally stable, eliminating the CFL constraint imposed by the high oxygen diffusion coefficient and allowing the same Δt_diff for all substrates regardless of D_k.

### Cell Agent Model

Cells are discrete agents with identity (cell_id, parent_id, generation), position (*x, y*), morphology (diameter, shape path, orientation), cell-cycle phase ∈ {*G*_0_, *G*_1_, *S, G*_2_, *M*}, ecDNA state (count and cargo), sampled local environment (O_2_, glucose), derived flags (is_hypoxic, is_quiescent), and effective behavioral rates. Nine cell types are modelled, with parameters summarized in Table 3.

**Table 3.** Cell types and behavioral parameters used in the 6mm publication configuration. Astrocyte division time corresponds to the slow turnover observed in adult cortex; tumor division time uses the 24-hour value calibrated for GBM. Apoptosis rates are 10^−4^/hr for normal lineages and 5 × 10^−5^/hr for tumor cells (further reduced under ecDNA-driven survival signaling).

| Cell type | Can<br>divide | Division<br>time (hr) | Migration<br>speed ($\mu\text{m}/\text{hr}$) | Key behaviors |
| --- | --- | --- | --- | --- |
| Neuron | no | — | 0 | Post-mitotic, high $\text{O}_2$ demand |
| Astrocyte | yes | $168 \pm 24$ | 3 | Contact-inhibited, reactive transition |
| Oligodendrocyte | no | — | 1 | Sensitive to hypoxia |
| Microglia | no | — | 30 | Resident immune, chemokine-activated |
| Endothelial | yes | $60 \pm 12$ | 10 | VEGF-responsive, vessel sprouting |
| Pericyte | no | — | 5 | Stabilizes endothelial vessels |
| Tumor | yes | $24 \pm 8$ | 10 | Non-contact-inhibited, ecDNA-bearing |
| Recruited<br>Immune | yes | $36 \pm 8$ | 35 | Chemotaxis, exhaustion dynamics |
| Necrotic | no | — | 0 | Lysing debris, blocks space |

#### Proliferation

Each hour, dividing-capable cells advance their cycle clock by Δ*t* provided the resource gate passes:

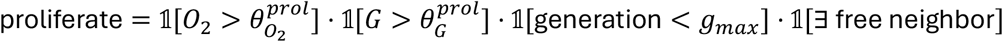

Phase durations are fractions of the cell-type-specific total division time *T*: 0.40*T* (G_1_), 0.30*T* (S), 0.15*T* (G_2_), 0.15*T* (M). If resources drop below threshold during S/G_2_/M, the cell arrests (clock pauses, phase preserved). On M-phase completion with an empty adjacent voxel available, a daughter is placed at that voxel, both cells reset to G_1_, and generation is incremented.

#### Migration

Motile cells compute a velocity vector as a weighted sum of persistence, chemotaxis, and noise:

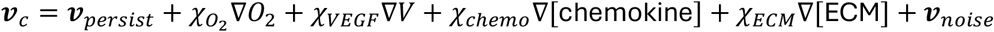

where ***v***_*persist*_ = ***v***_*prev*_ ⋅ exp(−Δ*t*/*τ*_*pers*_) provides directional memory and the χ coefficients are cell-type-specific chemotaxis sensitivities. Displacement over Δ*t* is (speed ⋅ cosθ,speed ⋅ sinθ), rounded to integer micrometers. Volume exclusion rejects moves that would place the cell within one cell-diameter of another; rejected moves retry an alternate direction or hold position.

#### Cell death

Three pathways: (i) necrosis triggers when 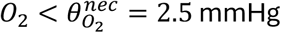 for *τ*_*nec*_ = 6 consecutive hours, converting the cell to a necrotic carcass that blocks space and is lysed stochastically at rate 0.01/hr; (ii) apoptosis fires each hour with *P*_*apo*_ = *a*_*eff*_ ⋅ Δ*t* and removes the cell immediately; (iii) immune kill fires when a recruited immune cell or activated microglia is within *r*_*kill*_ = 15 μm of a tumor cell, with kill probability 0._2_5 per maturation event (2.5 hr synapse maturation), capped at three kills per immune cell before exhaustion. The hypoxia-sensitive cell types (tumor, neuron, oligodendrocyte) carry a hypoxia-apoptosis multiplier of 50× to accelerate death under prolonged severe hypoxia.

#### Angiogenesis

When local *VEGF* exceeds θ_V = 5 nM adjacent to an endothelial cell, sprouting may occur. With probability 0.01/hr the tip cell extends one voxel up the *VEGF* gradient, a stalk cell occupies the vacated position, and the new vessel’s vascular density rises to its mature value over τ_mat = 48 hr. Anastomosis with another vessel completes a loop and activates the new path as an O_2_ and glucose source.

#### Immune module

The immune component simulates GBM-realistic immunosuppression. Each immunological synapse requires 2.5 hr to mature and carries 25% kill probability per maturation, substantially below the ∼70% typical of generic immune models, reflecting the GBM immunosuppressive microenvironment. T cells become exhausted after approximately three kills. Activation requires high local tumor density. Recruitment is restricted to 0.3%/hr to reflect limited infiltration across the blood-brain barrier. Together these parameters produce tumor growth despite immune pressure, matching the clinical reality of GBM immune evasion.

### ecDNA Replication and Segregation

ecDNA replicates during S phase and segregates during M phase. We model these as two sequential stochastic steps so that both replication fidelity and partition randomness contribute to within-lineage heterogeneity.

Replication (S phase). Given parent copy count N at S-phase entry, the replicated pool N’ is drawn as N original copies plus N independent Bernoulli replications with success probability p_rep = 0.95:

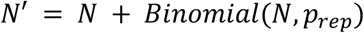

This yields a mean replication ratio of 1 + p_rep ≈ 1.95, slightly below the perfect doubling chromosomal DNA achieves, consistent with reported per-copy replication efficiency for circular ecDNA.

Segregation (M phase). The replicated pool N’ is partitioned between two daughter cells by independent Bernoulli trials with p_seg = 0.5:

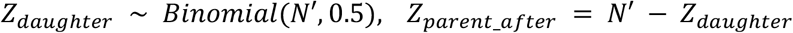

Conservation Z_daughter + Z_parent_after = N’ holds at the division event. Copy number is clamped to [0, Z_max] with Z_max = 100. The segregation depends only on N’ and p_seg; it does not depend on the local microenvironment U, which is the property that makes ecDNA a valid instrument by construction in the simulator.

### *EGFR* Expression and Hypoxia Coupling

*EGFR* protein per cell is computed as a function of ecDNA copy number and local hypoxia status, the latter modeling HIF-2α-driven translational upregulation of *EGFR* under hypoxia (Franovic et al. 2007^34^):

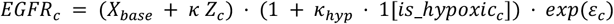

with X_base = 2.89 (baseline expression from chromosomal copy), κ = 1.21 (per-copy contribution from ecDNA-encoded *EGFR*), κ_hyp = 1.5 (default hypoxia-*EGFR* coupling, producing 2.5-fold upregulation in hypoxic cells), and ε_c ∼ N(0, 0.1^2^) (log-normal transcriptional noise).

The κ_hyp parameter is the hypoxia-*EGFR* coupling strength varied in the multi-scenario robustness analysis: κ_hyp ∈ {1.5, 0.5, 0.0} corresponding to baseline / reduced / removed conditions. Setting κ_hyp = 0 severs the Hypoxia → *EGFR* edge in the SCM and is the cleanest test of whether observed OLS bias is attributable to the modeled confounding pathway.

The downstream effects of ecDNA copy number on cellular phenotypes are the ground-truth causal effects against which IV and discovery methods are benchmarked:

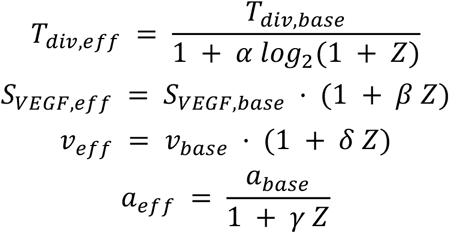

with default ground-truth coefficients α = 0.30 (per-copy effect on division rate), β = 0.10 (*VEGF* amplification), δ = 0.05 (migration), γ = 0.50 (apoptosis suppression). These coefficients are written to a per-run at simulation start and are treated as the known ground truth against which estimators are scored.

### Initialization and Equilibration

Normal tissue is populated at 10^−3^ cells/µm^2^ with cell-type fractions calibrated to adult cortical gray matter: 45% neurons, 25% astrocytes, 12% oligodendrocytes, 7% microglia, 4% endothelial, 2% pericytes. The vascular network is placed first as a branching capillary pattern with inter-capillary spacing of ∼150 µm; pericytes are placed adjacent to endothelial cells. Tumor founders (1–3 cells with initial Z = 20 *EGFR*-bearing ecDNA copies) are placed near the domain center.

Before the recorded time series begins, environment fields are equilibrated for a burn-in period (t_burn = 30 hr for 2 mm runs, 40 hr for 6 mm runs) during which the LOD diffusion solver runs but no cell behaviors execute. This ensures the recorded t = 0 state reflects steady-state O_2_ and glucose distributions consistent with the placed vasculature rather than the uniform initial condition.

### Main Simulation Loop

For each main step *t* = 1,_2_, …, *T*_*total*_:

1. **Phase 1: Environment diffusion**. For each substrate, run ⌊Δ*t*/Δ*t*_*diff*_⌋ = 100 implicit-LOD sub-steps (computing sources/sinks, solving x-sweep then y-sweep, applying decay and clamps). Derived fields (pH from lactate, hypoxia flags from O_2_) are updated at the end of Phase 1.
2. **Phase 2: Cell behavior** (cells processed in randomly shuffled order). For each living cell: sample local environment via bilinear interpolation, update hypoxic/quiescent flags, check necrosis and apoptosis, evaluate state transitions, advance the cell cycle (including division with binomial ecDNA segregation), and attempt migration (with volume exclusion).
3. **Phase 3: Angiogenesis**. For each endothelial cell with adjacent *VEGF* above threshold, attempt sprouting.
4. **Phase 4 : Immune recruitment**. For each vascular position with chemokine above threshold, stochastically spawn a recruited immune cell.

Per-hour outputs serve two purposes: (i) post-hoc IV and discovery analyses operate on the final-timestep cells file, and (ii) the lineage file provides every division event needed to validate the binomial segregation assumption empirically.

### Output Analysis G Validation

Using simulation output, we estimate the ecDNA→phenotype effects through three approaches: OLS regression, which is biased by confounding; two-stage least squares using ecDNA as the instrument, which recovers the true configured values; and sibling comparison, which controls all shared confounders. These targets differ in what the data can identify, and we are explicit about this: β (*VEGF*) and δ (migration) are continuous per-cell phenotypes recorded at every snapshot, so 2SLS recovers them directly on the structural scale in every scenario; α (division) is a dynamic rate, point-identified from inter-division intervals (with the configured base cycle time held fixed as a known constant) only in the unconfounded κ_hyp = 0 runs, where *EGFR* is an exact function of ecDNA; and γ (survival) is not identified, because the configured baseline apoptosis rate is so low (5×10^−5^/hr) that essentially no tumor cell dies and there is no survival selection for γ to act on — we therefore treat γ as a configured design parameter rather than an estimation target. We evaluate each identified estimator using relative error (the absolute difference between estimated and true values normalized by the true value), coverage (whether the 95% confidence interval includes the true value), and the bias ratio of the OLS estimate to the IV estimate, where values greater than one indicate positive confounding.

We validate the segregation mechanism by verifying that daughter ecDNA fractions follow a Binomial(N, 0.5) distribution, testing that the mean fraction equals 0.5 by t-test and that the variance matches the expected p(1-p)/N. Instrument strength is confirmed by requiring that the first-stage F-statistic exceeds 10 for all analyses.

### Structural Causal Model for ecDNA-Driven Tumor Biology

We formalize the causal structure of ecDNA-driven tumor phenotypes. An SCM consists of a set of endogenous variables V, a set of exogenous variables U, and a set of structural equations F that determine each endogenous variable as a function of its parents and an exogenous noise term. The graphical representation of the SCM is a directed acyclic graph (DAG) where edges encode direct causal relationships. Let the endogenous variables be:

- **Z** = ecDNA copy number (instrument, range 0-100)
- X = *EGFR* expression level (exposure, normalized units)
- Y_**1**_ = cell cycle duration, i.e., time to division (outcome)
- Y_**2**_ = migration speed (outcome, μm/hour)
- Y_**3**_ = *VEGF* secretion rate (outcome, amol/hour)
- Y_**4**_ = apoptosis rate (outcome, probability/hour)
- M = hypoxia status (mediator/confounder, binary)

Let the exogenous variables be:

- U_**O2**_ = local oxygen concentration (mmHg), determined by vascular distance and consumption
- U_**gluc**_ = local glucose concentration (mM)
- ε_**X**_, **ε**_**Y1**_, …, **ε**_**Y4**_ = independent noise terms representing biological variability

### Structural Equations

The causal mechanisms are specified by the following structural equations describing gene dosage, hypoxia status, cell-cycle duration, migration speed, among others.

Gene dosage (Z → X) was calculated where *X*_base_ = 1.0 is baseline *EGFR* expression from the chromosomal copy, *κ* = 1.2 is the expression contribution per ecDNA copy, *Z*_max_ = 100 caps the copy number effect, and *σ*_*X*_ = 0.1 introduces cell-to-cell variability in transcription/translation efficiency:

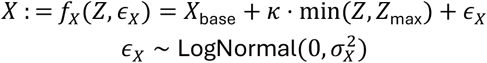

Hypoxia status (U_O_2_ → M) was calculated where *τ*_*hypoxia*_ = 18 mmHg is the threshold for HIF-2α stabilization. This is a deterministic threshold function reflecting the sharp oxygen dependence of HIF degradation.

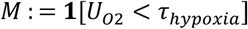

Cell cycle duration (X, M, U_O2 → Y_1_) was calculated where *T*_base_ = 24 hours is the baseline division time, *α* = 0.3 is the *EGFR*-mediated acceleration, and *τ*_*prolif*_ = 8 mmHg is the minimum oxygen for proliferation. The indicator functions enforce the biological constraint that severely hypoxic cells arrest proliferation:

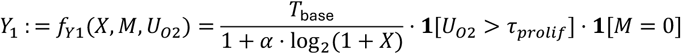

Migration speed (X, M → Y_2_) was calculated where *v*_base_ = 10 μm/hour is baseline migration speed, *δ* = 0.05 is the *EGFR*-mediated migration boost, and *ψ* = 2.0 is the hypoxia-induced “Go” switch fors. This encodes the “Go or Grow” hypothesis: hypoxic cells upregulate migration while suppressing proliferation:

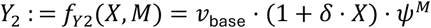

*VEGF* secretion (X, M → Y_3_) was calculated where *S*_base_ = 600 amol/hour is baseline secretion, *β* = 0.1 is *EGFR*-mediated amplification (square-root form reflecting receptor saturation), and the final term captures the 5-fold upregulation of *VEGF* under hypoxia via HIF-2α.

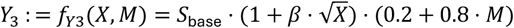

Apoptosis rate (X → Y_4_) was calculated where *a*_base_ = 5 × 10^−5^ per hour is baseline apoptosis rate and *γ* = 0.5 is the *EGFR*-mediated survival effect. High *EGFR* activates PI3K/AKT survival signaling, reducing apoptosis.

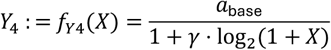

### The Confounding Structure

The critical feature of this SCM is that oxygen (U_O2) creates confounding between *EGFR* expression and phenotypes through two pathways: **U_O2 → M → Y**: Hypoxia directly affects phenotypes via the Go-or-Grow switch and HIF-mediated transcription; **U_O2 → (cell selection) → observed X**: Cells in hypoxic regions may have systematically different ecDNA distributions due to differential survival or proliferation However, the *segregation* of ecDNA at cell division is independent of oxygen, where 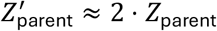 after S-phase replication. This conditional independence is the source of instrument validity. Here, the causal effect of X on any outcome Y_i is identified by the IV estimand:

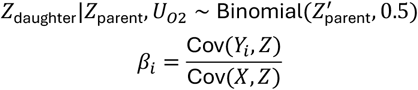

This is valid because: (1) Z affects X (relevance: gene dosage), (2) Z ⊥⊥ U_O2 (independence: random segregation), and (3) Z affects Y only through X (exclusion: ecDNA acts via gene expression).

### Two-Stage Least Squares Estimation

Given the causal structure above, we recover the effect of X on Y using two-stage least squares (2SLS): In the first stage, we regress exposure on instrument and covariates, where W represents observed covariates (e.g., glucose, spatial position). The predicted values 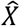 capture only the variation in X attributable to the random instrument Z. In the second stage, we regress outcome on predicted exposure, wheretThe coefficient *β*_1_ is the causal effect of X on Y, purged of confounding by U because 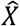 contains only variation from the randomized component (ecDNA segregation).

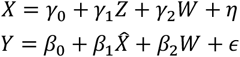

The IV estimator is the ratio of the reduced-form effect (Z→Y) to the first-stage effect (Z→X), isolating the X→Y pathway and the Instrument Strength Diagnostics. A weak instrument (one that barely affects X) produces biased and imprecise IV estimates. We assess instrument strength via the first-stage F-statistic:

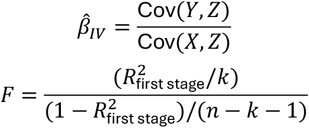

where k is the number of instruments and n is the sample size. We require F > 10 to ensure reliable inference. For ecDNA, we expect strong instruments because gene dosage effects are large: each additional ecDNA copy increases expression by approximately *κ* = 0.5 units, and copy numbers range from 0 to 100+.

### Sibling Comparison Design

An alternative identification strategy exploits the within-family randomization of ecDNA segregation. When a parent cell with N ecDNA copies divides: Daughter A receives *Z*_*A*_ ∼ Binomial(*N*^′^, 0.5) copies; Daughter B receives *Z*_*B*_ = *N*^′^ − *Z*_*A*_ copies, where *N*^′^ ≈ _2_*N* accounts for ecDNA replication during S phase. Because siblings share identical genetics, spatial origin, and temporal history, any difference in their phenotypes must be caused by the random difference in their ecDNA counts, where Δ*X* = *X*_*A*_ − *X*_*B*_ arises solely from Δ*Z* = *Z*_*A*_ − *Z*_*B*_. This design eliminates all confounders shared between siblings, including unobserved ones:

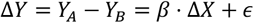

### Sensitivity Analysis

We assessed robustness to exclusion-restriction violations using three complementary tools.

#### Rosenbaum bounds

We computed the critical odds ratio Γ at which the IV estimate loses statistical significance at α = 0.05. Γ is the maximum departure from random treatment assignment a hidden binary confounder could induce while still permitting the observed association. Critical Γ values were obtained by numerical search; we report the smallest Γ at which the upper bound of the one-sided p-value exceeds 0.05.

#### E-values

The E-value for a risk ratio RR is the minimum strength, on the risk-ratio scale, that an unmeasured confounder would need with both the instrument and the outcome to fully account for the observed effect:

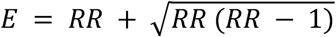

with the analogous expression evaluated at the 95% confidence-interval bound to give a confidence-interval E-value.

#### Placebo (negative-control) outcomes

We tested whether ecDNA copy number predicts variables that should have no causal connection under the SCM. The three negative-control outcomes were cell x-coordinate, cell y-coordinate, and distance to tumor center. Null associations are consistent with the construction of the simulator and provide a sanity check on the analysis pipeline.

### Causal Discovery

To complement effect estimation, we applied the PC constraint-based causal discovery algorithm to recover the underlying DAG (directed acyclic graph) from data without specifying which edges to test. The algorithm begins from a complete undirected graph and tests conditional independence between each pair of variables given conditioning sets of increasing size, removing edges where conditional independence holds. Remaining edges are oriented using collider detection and acyclicity constraints.

Significance level for the independence tests was α = 0.05 using the Fisher Z-test for partial correlations; maximum conditioning set size was k = 3. Variables included were Z (ecDNA), X (*EGFR*), M (hypoxia), Y_1_ (proliferation), Y_2_ (migration), Y_3_ (*VEGF*), and Y_4_ (survival). Glucose was excluded from the variable set because it is highly correlated with O_2_ (both determined by vascular distance); including redundant variables substantially degraded discovery performance in pilot runs.

Recovered graphs were compared to the ground-truth DAG using Structural Hamming Distance, precision, recall, and F1 score.

### Experimental Design

#### Multi-Scenario Factorial Analysis

To assess SIV-framework behavior across confounder strengths and tumor scales, we ran a 2 × 3 × 5 factorial design: two spatial scales (2 mm and 6 mm), three hypoxia-*EGFR* coupling scenarios (κ_hyp = 1.5 baseline, 0.5 reduced, 0.0 removed), and five independent random seeds per (scale × scenario) condition, for 30 simulations total. All reported effect estimates carry seed-to-seed standard deviations across the five seeds.

#### Time-resolved analysis

To characterize how confounder visibility and effect-estimate stability evolve as a tumor matures, IV and PC analyses were re-run at five intermediate timepoints per simulation: t = 80, 120, 160, 200, and 240 hr for the 2 mm runs; t = 100, 150, 200, 250, and 300 hr for the 6 mm runs. The cells file from each timepoint was treated as an independent cross-section for the analysis.

#### Spatial Heterogeneity Analysis

Tumor cells were stratified into three concentric zones by distance from the tumor-seed centroid (core, margin, and infiltrating) and the migration effect δ was estimated separately within each zone by structural 2SLS using ecDNA copy number as the instrument. We ran this analysis on two configurations: (i) the standard constant-δ simulator (negative control), and (ii) a modified simulator in which δ was set to a known spatial gradient via the optional ecDNA_migration_spatial mode (core < 780 µm: δ = 0.03; margin 780–970 µm: δ = 0.07; infiltrating > 970 µm: δ = 0.05), to test whether the per-zone estimator recovers a planted gradient.

### Validation Metrics

For each estimator (OLS, 2SLS, sibling comparison), we report three diagnostics:

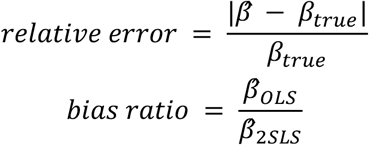

plus 95% CI coverage of the true value. The segregation rule was empirically validated by testing that the mean daughter fraction equals 0.5 (one-sample t-test) and that observed variance matches the binomial expectation:

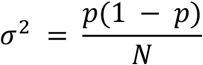

Instrument strength was confirmed by requiring first-stage F > 10 in every analysis.

## Code availability

https://github.com/davcraig75/causanta.

## Data availability

Simulation parameters and outputs available as supplementary material.

## Notes

https://github.com/davcraig75/causanta

### Competing Interest Statement

The authors have declared no competing interest.

