## Supplemental Materials for "Extrachromosomal DNA as a Causal Instrument for Spatial Multi-Omics"

### CAUSANTA Supplementary Materials

#### Statistical Methods for Nature Methods Publication

This document provides detailed methodological specifications for the CAUSANTA framework, suitable for peer review and reproducibility.

---

#### Plain-language reading guide

The main manuscript describes *what* we did and *what we found*; this supplement describes the *mathematical machinery* underneath each result so that a statistician or computational biologist can reproduce it from scratch. For a biological reader who wants to understand a particular result without going through the full derivation, the relevant sections map onto the main-text claims as follows.

If you want to know how we **estimate causal effects** (the OLS-vs-IV comparison in §Causal Effect Recovery): start with §2.1 (Two-Stage Least Squares — the actual two-step regression we run) and then §2.3 (Instrument Strength Diagnostics — what the F-statistic is and what threshold matters). §2.2 (Standard Errors) explains why the reported error bars are larger than you might expect from the point estimates alone.

If you want to know how we **test robustness** (the Rosenbaum bounds and E-values in §Sensitivity Analysis): jump to §6. The intuition is the same one the main text uses — $\Gamma^*$ is the critical confounder strength that would overturn the result, E-value is the same idea on a risk-ratio scale, and placebo tests check that the instrument fails on outcomes it should not predict. The math is here.

If you want to know how we **score graph recovery** (the F1 scores in §Causal Discovery): §7 specifies the PC and GES algorithms with their parameter choices (α, BIC penalty), how we compute precision/recall/F1, and how the bootstrap stability test (§7.3) handles seed-to-seed variation.

Sections 3 (Bootstrap CIs), 4 (Power Analysis), and 5 (Heterogeneity) cover three things that come up only briefly in the main text but matter for any future application to real data — how we get confidence intervals when the IV estimator's sampling distribution is non-Gaussian, how to compute the sample size needed for a target effect size, and how we stratify estimates spatially across tumor regions (the Cochran's Q and $I^2$ statistics).

Section 8 lists every numerical parameter used in the canonical 6 mm publication run so that the entire dataset and analysis is reproducible from the configuration file alone.

---

### Table of Contents


1. Mathematical Framework
2. Instrumental Variable Estimation
3. Bootstrap Confidence Intervals
4. Power Analysis
5. Heterogeneity Analysis
6. Sensitivity Analysis
7. Causal Discovery
8. Simulation Parameters
9. Success Criteria
10. Software Implementation


---

### 1. Mathematical Framework

#### 1.1 Structural Causal Model

The CAUSANTA simulator implements the following structural equations:

**ecDNA Segregation (Instrument Generation)**

```
N_daughter ~ Binomial(N_parent, 0.5)
```

**Gene Expression (First Stage; gene-dosage + HIF-2α coupling)**

```
EGFR_i = [base + κ · ecDNA_i] · [1 + κ_hyp · is_hypoxic_i] · exp(ε_i)
       base = 2.89, κ = 1.21, κ_hyp ∈ {1.5, 0.5, 0.0}
(baseline/reduced/removed),
       ε_i ~ N(0, 0.1²)
```

**Phenotypic Effects (Structural Equations; ecDNA → phenotype via EGFR)**

```
T_div_eff_i = T_base / (1 + α · log2(1 + ecDNA_i))
VEGF_i      = VEGF_base · (1 + β · ecDNA_i) · 1[is_hypoxic_i]   (gated
secretion)
Migration_i = v_base · (1 + δ · ecDNA_i)
Apoptosis_i = a_base / (1 + γ · ecDNA_i)
```

**Confounding Structure (matches simulator exactly)**

```
is_hypoxic_i (binary HIF threshold indicator) → EGFR_i      via HIF-2α
translation
is_hypoxic_i → VEGF_i                                       via hypoxia-gated
secretion
O2_local_i (continuous; < 18 mmHg ⇒ is_hypoxic_i = 1)
```

The simulator's coupling to oxygen is threshold-mediated (step function at 18 mmHg), so the structural causal signal lives in the binary is_hypoxic indicator, not the continuous

`O2_local` field. Causal discovery in this paper uses `is_hypoxic` as the confounder variable; with continuous O2 in the variable set, late 6 mm runs give only F1 ≈ 0.76 because every tumor cell is already above the threshold for the effect.

#### 1.2 Ground Truth Parameters

| Symbol | Parameter | Default Value | Biological Interpretation |
|---|---|---|---|
| α | ecDNA_effect_on_division | 0.30 | 30% faster division per doubling of ecDNA |
| β | ecDNA_effect_on_VEGF | 0.10 | 10% more VEGF per ecDNA copy |
| δ | ecDNA_effect_on_migration | 0.05 | 5% faster migration per ecDNA copy |
| γ | ecDNA_effect_on_survival | 0.50 | 50% survival boost per ecDNA copy |
| κ | EGFR per ecDNA copy | 1.21 | Per-copy gene-dosage coefficient on EGFR |
| κ_hyp | hypoxia EGFR upregulation | 1.5 (baseline) | HIF-2α multiplier; swept {1.5, 0.5, 0.0} |

#### 1.3 Causal DAG

```
                          is_hypoxic (binary HIF indicator)
                               |
                    ┌──────────┴──────────────┐
                    │ HIF-2α (κ_hyp)           │ hypoxia gating
                    ▼                          ▼
ecDNA_count (Z) ──gene dosage (κ)──→ EGFR (D) ──→ VEGF, migration, division,
survival (Y)
        ▲
        |
Binomial(N, 0.5)
(random segregation; the instrument)
```

**Key Properties:** - $Z \perp U$ : ecDNA segregation is independent of confounders (random mechanism) - $Z \to D$ : ecDNA causally affects EGFR expression (gene dosage, κ = 1.21 per copy) - is_hypoxic → D : HIF-2α-mediated translational upregulation under hypoxia (confounder edge; strength controlled by κ_hyp) - is_hypoxic → Y_VEGF : direct hypoxia-gated VEGF secretion (independent of D) - $Z \perp\!\!\!\perp Y \mid D$ : ecDNA affects outcomes only through EGFR (exclusion restriction)

When κ_hyp = 0 the (is_hypoxic → D) edge is severed; OLS and IV converge to within ±1% at both 2 mm and 6 mm scales (Table 2 of the main manuscript).

### 2. Instrumental Variable Estimation

#### 2.1 Two-Stage Least Squares (2SLS)

**Stage 1 (First Stage):**

$$D_i = \pi_0 + \pi_1 * Z_i + \pi_2 * X_i + \eta_i$$
$$\hat{D}_i = \hat{\pi}_0 + \hat{\pi}_1 * Z_i + \hat{\pi}_2 * X_i$$

**Stage 2 (Second Stage):**

$$Y_i = \beta_0 + \beta_1 * \hat{D}_i + \beta_2 * X_i + \varepsilon_i$$

Where: - $Z_i$ = ecDNA copy number (instrument) - $D_i$ = EGFR expression (endogenous treatment) - $Y_i$ = Phenotypic outcome - $X_i$ = Observed covariates (optional)

#### 2.2 Standard Error Correction

The naive second-stage standard errors are incorrect. We use the corrected formula:

$$SE(\hat{\beta}_{IV}) = \text{sqrt}(\sigma^2 * (\hat{X}'\hat{X})^{(-1)})$$

Where $\sigma^2$ is estimated from residuals using original D (not $\hat{D}$):

$$\sigma^2 = \Sigma(Y_i - \hat{\beta}_0 - \hat{\beta}_1 * D_i - \hat{\beta}_2 * X_i)^2 / (n - k)$$

#### 2.3 Instrument Strength Diagnostics

**F-Statistic:**

$$F = (R^2_{first} / k) / ((1 - R^2_{first}) / (n - k - 1))$$

| F-Statistic | Interpretation |
|---|---|
| $F > 10$ | Strong instrument (Staiger & Stock rule) |
| $F > 16.38$ | <5% maximal IV size distortion |
| $F > 8.96$ | <10% maximal IV size distortion |

**Partial $R^2$:**

$$R^2_{partial} = (R^2_{full} - R^2_{restricted}) / (1 - R^2_{restricted})$$

#### 2.4 Wu-Hausman Endogeneity Test

Tests whether OLS and IV estimates differ significantly.

**Augmented Regression:**

$$Y_i = \beta_0 + \beta_1 * D_i + \beta_2 * X_i + \theta * \hat{v}_i + \varepsilon_i$$

Where $\hat{v}_i = D_i - \hat{D}_i$ (first-stage residuals).

**Test:** H0: $\theta = 0$ (no endogeneity) - If $p < 0.05$, OLS is biased → use IV

##### 2.5 Anderson-Rubin Confidence Intervals

Robust to weak instruments:

$$AR(\beta_0) = n * (SSR_{restricted}(\beta_0) - SSR_{unrestricted}) / SSR_{unrestricted}$$

Under H0: $\beta = \beta_0$, $AR \sim \chi^2(1)$

The AR confidence set inverts this test:

$$CI_{AR} = \{\beta_0 : AR(\beta_0) \leq \chi^2_{1,\alpha}\}$$

---

#### 3. Bootstrap Confidence Intervals

##### 3.1 BCa (Bias-Corrected and Accelerated) Bootstrap

The BCa method adjusts for bias and skewness in the bootstrap distribution.

**Algorithm:** 1. Compute $\hat{\theta}$ from original sample 2. Draw B bootstrap samples, compute $\theta^*_b$ for each 3. Compute bias correction: $\hat{z}_0 = \Phi^{-1}(\#\{\theta^*_b < \hat{\theta}\} / B)$ 4. Compute acceleration (jackknife influence): $\hat{a} = \Sigma(\hat{\theta}_{(-i)} - \hat{\theta}_{(\cdot)})^3 / (6 * [\Sigma(\hat{\theta}_{(-i)} - \hat{\theta}_{(\cdot)})^2]^{3/2})$ 5. Adjusted percentiles: $\alpha_1 = \Phi(\hat{z}_0 + (\hat{z}_0 + z_{\alpha/2}) / (1 - \hat{a}(\hat{z}_0 + z_{\alpha/2})))$ $\alpha_2 = \Phi(\hat{z}_0 + (\hat{z}_0 + z_{1-\alpha/2}) / (1 - \hat{a}(\hat{z}_0 + z_{1-\alpha/2})))$ 6. CI = $[\theta^*_{(\alpha_1)}, \theta^*_{(\alpha_2)}]$

##### 3.2 Bootstrap for IV Estimates

Each bootstrap replicate: 1. Resample $(Z_i, D_i, Y_i)$ with replacement 2. Run full 2SLS procedure 3. Store IV coefficient

##### 3.3 Testing Segregation Randomness

**Chi-Square Test:**

$$\chi^2 = \Sigma (O_k - E_k)^2 / E_k$$

Where: - $O_k$ = Observed count in bin k - $E_k = n * P(daughter\_fraction \in bin_k \mid Binomial)$

**Kolmogorov-Smirnov Test:**

$$D = \max|F_n(x) - F_0(x)|$$

Where $F_0$ is the theoretical CDF under Binomial(N, 0.5).

---

### 4. Power Analysis

#### 4.1 Analytical Power for 2SLS

**Asymptotic Distribution:**

$$\sqrt{n}(\hat{\beta}_{IV} - \beta) \to_d N(0, \sigma^2 / (\pi_1^2 * Var(Z)))$$

**Standard Error of IV:**

$$SE_{IV} \approx sqrt(\sigma^2 / (n * R^2_{first} * Var(Z)))$$

**Power:**

$$Power = 1 - \Phi(z_{1-\alpha/2} - |\beta| / SE_{IV}) + \Phi(-z_{1-\alpha/2} - |\beta| / SE_{IV})$$

#### 4.2 Required Sample Size

For target power (1 - β) at significance level α:

$$n \approx (z_{1-\alpha/2} + z_{1-\beta})^2 * \sigma^2 / (\beta^2 * R^2_{first} * Var(Z))$$

#### 4.3 Minimum Detectable Effect (MDE)

Given sample size n and target power:

$$MDE = (z_{1-\alpha/2} + z_{1-\beta}) * SE_{IV}$$

#### 4.4 Power Curve Parameters

| Effect Size | Required n (80% power) | Required n (90% power) |
|---|---|---|
| 0.01 | ~10,000 | ~15,000 |
| 0.02 | ~2,500 | ~3,750 |
| 0.05 | ~400 | ~600 |
| 0.10 | ~100 | ~150 |
| 0.20 | ~25 | ~40 |

*(Assuming $R^2\_first = 0.884$, $\alpha = 0.05$)*

---

### 5. Heterogeneity Analysis

#### 5.1 Stratified IV Estimation

Run separate IV analyses by stratum:

1. **By Region:**
   - Core: cells within 100μm of tumor center

  - Margin: cells 100-300μm from center
  - Infiltrating: cells >300μm from center
2. **By Hypoxia:**
  - Normoxic: $O2 > 20$ mmHg
  - Mildly hypoxic: 10-20 mmHg
  - Severely hypoxic: <10 mmHg
3. **By ecDNA Burden:**
  - Low: ecDNA < 10
  - Medium: 10-30
  - High: >30

##### 5.2 Cochran's Q Test for Heterogeneity

$$Q = \Sigma\, w\_k\, (\hat{\beta}\_k - \hat{\beta}\_pooled)^2$$

Where $w\_k = 1 / SE^2\_k$

Under H0 (homogeneity): $Q \sim \chi^2(K-1)$

##### 5.3 $I^2$ Statistic

$$I^2 = \max(0, (Q - (K-1)) / Q)$$

| $I^2$ | Heterogeneity Level |
|---|---|
| 0-25% | Low |
| 25-50% | Moderate |
| 50-75% | Substantial |
| >75% | Considerable |

##### 5.4 Radial Effect Profiles

Estimate effects as smooth function of distance from tumor center:

$$\beta(r) = f(r) \text{ estimated via local regression}$$

Kernel bandwidth selected by cross-validation.

---

#### 6. Sensitivity Analysis

##### 6.1 Rosenbaum Bounds

Assess sensitivity to unobserved confounding.

For treatment effect Γ (odds ratio of hidden bias):

$$P(p\_upper > \alpha \mid \Gamma) = \text{robustness to confounding at level } \Gamma$$

**Interpretation:** - Γ = 1: No hidden bias (randomization) - Γ = 2: Unobserved confounder doubles treatment odds - Critical Γ*: Smallest Γ where p_upper > α

##### 6.2 E-Value

Minimum confounding strength to explain away observed effect:

```
E-value = RR + sqrt(RR * (RR - 1))
```

Where RR is the risk ratio estimate.

**Interpretation:** - E-value = 3: Confounder must have RR ≥ 3 with both treatment and outcome to explain away effect - Higher E-value → more robust to confounding

##### 6.3 Instrument Exclusion Tests

**Falsification Tests:** 1. Test Z → pre-treatment outcomes (should be null) 2. Test Z → known non-pathway outcomes (should be null) 3. Overidentification test (if multiple instruments)

---

#### 7. Causal Discovery

##### 7.1 PC Algorithm

Constraint-based causal discovery:

1. Start with complete undirected graph
2. Remove edges based on conditional independence tests
3. Orient edges using v-structures and propagation rules

**Independence Test:** Partial correlation test with Fisher's z-transformation:

```
z = 0.5 * log((1+r)/(1-r)) * sqrt(n-k-3)
```

##### 7.2 GES (Greedy Equivalence Search)

Score-based causal discovery:

1. Forward phase: Add edges that improve BIC score
2. Backward phase: Remove edges that improve BIC score
3. Return equivalence class

**BIC Score:**

```
BIC = -2 * log L + k * log(n)
```

##### 7.3 Bootstrap Stability

Edge stability across bootstrap samples:

```
Stability(edge) = #{bootstrap samples with edge} / B
```

Threshold: edges with stability > 0.5 are retained.

---

#### 8. Simulation Parameters

Default values below are taken from the family of parameter files in `causanta/simulate/params/` used in the multi-scale reliability sweep:

| File family | Scale | Total hr | Burn-in | n seeds | Purpose |
|---|---|---|---|---|---|
| `robustness_{baseline,reduced,removed}.json` | 1 mm | 160 | 20 | 1 (42) | small-domain calibration |
| `large_{baseline,reduced,removed}.json` | 2 mm | 240 | 30 | 1 (42) | mid-scale single seed |
| `xlarge_{baseline,reduced,removed}.json` | 6 mm | 300 | 40 | 1 (42) | publication-scale single seed |
| `multiseed_2mm_{baseline,reduced,removed}_seed{43-46}.json` | 2 mm | 240 | 30 | 4 | 2 mm reliability bounds (n = 5 with seed 42) |
| `multiseed_6mm_{baseline,reduced,removed}_seed{43-46}.json` | 6 mm | 300 | 40 | 4 | 6 mm reliability bounds (n = 5 with seed 42) |

The `egfr_hypoxia_upregulation` parameter ($\kappa$_hyp) takes values 1.5 / 0.5 / 0.0 for the baseline / reduced / removed scenarios respectively.

##### 8.1 Domain Configuration

| Parameter | 1 mm | 2 mm | 6 mm | Description |
|---|---|---|---|---|
| width_um | 1000 | 2000 | 6000 | Domain width (μm) |
| height_um | 1000 | 2000 | 6000 | Domain height (μm) |
| env_grid_um | 10 | 10 | 10 | Environment grid spacing (μm) |

##### 8.2 Time Configuration

| Parameter | 1 mm | 2 mm | 6 mm | Description |
|---|---|---|---|---|
| total_hours | 160 | 240 | 300 | Simulation duration (hr) |

| Parameter | 1 mm | 2 mm | 6 mm | Description |
|---|---|---|---|---|
| dt_diffusion_hr | 0.01 | 0.01 | 0.01 | Diffusion sub-step (36 s) |
| output_interval_hr | 1 | 1 | 1 | Output frequency (hr) |
| burnin_hours | 20 | 30 | 40 | Burn-in equilibration before t = 0 (hr) |

##### 8.3 Cell Type Parameters (canonical 6 mm publication configuration)

| Cell Type | Can Divide | Division Time (hr) | Apoptosis Rate (/hr) | Migration Speed (μm/hr) |
|---|---|---|---|---|
| Neuron | no | — | 1e-4 | 0 |
| Astrocyte | yes | 168 ± 24 | 1e-4 | 3 |
| Oligodendrocyte | no | — | 1e-4 | 1 |
| Microglia | no | — | 1e-4 | 30 |
| Endothelial | yes | 60 ± 12 | 1e-4 | 10 |
| Pericyte | no | — | 1e-4 | 5 |
| Tumor | yes | **24 ± 8** | 5e-5 | 10 |
| RecruitedImmune | yes | 36 ± 8 | 1e-3 | 35 |
| Necrotic | no | — | 0 | 0 |

(Tumor division time is 24 hr in the publication-scale runs to match the doubling time observed in GBM clinical samples. The legacy 36 hr value appears in `default.json` for backwards compatibility but is not used for any reported result.)

##### 8.4 ecDNA Parameters

| Parameter | Value | Description |
|---|---|---|
| Initial copies | 20 | Starting ecDNA in seed tumor |
| Segregation p | 0.5 | Binomial probability |
| α (effect on division) | 0.30 | $T\_eff = T\_base / (1 + \alpha \cdot \log_2(1 + ecDNA))$ |
| β (effect on VEGF) | 0.10 | $VEGF\_eff = VEGF\_base \times (1 + \beta \cdot ecDNA)$ |
| δ (effect on migration) | 0.05 | $v\_eff = v\_base \times (1 + \delta \cdot ecDNA)$ |
| γ (effect on survival) | 0.50 | $a\_eff = a\_base / (1 + \gamma \cdot ecDNA)$ |

#### 9. Success Criteria

##### 9.1 Publication-Ready Metrics

| Criterion | Target | Measurement |
|---|---|---|
| IV Accuracy | <15% bias | |
| CI Coverage | ≥90% | Fraction containing true value |
| Power | ≥80% | At $\delta = 0.05$, $n = 1000$ |
| Segregation Test | $p > 0.05$ | KS test vs Binomial |
| First-stage F | >10 | Standard threshold |
| E-value | >2.0 | Robustness to confounding |

##### 9.5 Multi-Seed Reliability Bounds (canonical run set, 30 simulations)

The values below are aggregated from `output/multiseed_full_results.json` (n = 5 seeds per (scale, scenario) cell). Discovery uses `is_hypoxic` with scenario-aware ground truth (5 edges for baseline / reduced; 4 edges for removed). The bias % column is (OLS − IV) / IV × 100. Full per-seed values are reproduced in the main manuscript's Table 2.

| Scale | Scenario | HIF (κ_hyp) | OLS β | IV β | OLS bias % | F1 |
|---|---|---|---|---|---|---|
| 2 mm | baseline | 1.5 | 4.199 ± 0.133 | 3.981 ± 0.098 | +5.5 ± 1.4 | 1.000 ± 0.000 |
| 2 mm | reduced | 0.5 | 5.611 ± 0.210 | 5.277 ± 0.162 | +6.3 ± 1.8 | 0.874 ± 0.118 |
| 2 mm | removed | 0.0 | 6.397 ± 0.150 | 6.455 ± 0.136 | −0.9 ± 0.3 | 0.889 ± 0.000 |
| 6 mm | baseline | 1.5 | 4.113 ± 0.128 | 3.642 ± 0.110 | +12.9 ± 0.5 | 1.000 ± 0.000 |
| 6 mm | reduced | 0.5 | 5.080 ± 0.072 | 4.943 ± 0.137 | +2.8 ± 1.7 | 0.909 ± 0.203 |
| 6 mm | removed | 0.0 | 5.965 ± 0.230 | 6.030 ± 0.217 | −1.1 ± 0.4 | 0.824 ± 0.089 |

First-stage F-statistics across the 30 sims range from 37,810 to 229,655. Time-resolved analysis at five intermediate timesteps per run (`output/multiseed_timeseries_results.json`, Figure 7 of the main manuscript) shows F1 increasing monotonically and OLS bias decreasing as the tumor matures.

##### 9.2 Validation Benchmarks

1. **Segregation Validation:**
   - Mean daughter fraction: 0.50 ± 0.02

    - Variance matches Binomial prediction
    - No trend with parent ecDNA count
2. **First Stage Strength:**
    - F > 100 (very strong by design)
    - $R^2 > 0.80$ for ecDNA → EGFR
3. **Effect Recovery:**
    - β (VEGF) and δ (migration) recovered within 2 SE by 2SLS in every scenario (per-cell phenotypes); α (division) recovered from inter-division intervals on the κ_hyp = 0 runs (0.323/0.307 vs 0.30); γ (survival) not identified — negligible apoptosis (5e-5/hr) leaves no survival selection
    - IV estimates unbiased, OLS estimates show confounding bias
4. **Heterogeneity Detection:**
    - Detect planted heterogeneity when effect varies by region
    - Q-test sensitive to true heterogeneity

---

#### 10. Software Implementation

##### 10.1 Module Structure

```
causanta/
├── simulate/
│   ├── core.py          # Main simulation loop
│   ├── ecdna.py         # ecDNA segregation model
│   ├── sweep.py         # Parameter sweep system
│   └── params/          # Configuration files
├── analyze/
│   ├── iv.py            # 2SLS estimation + diagnostics
│   ├── bootstrap.py     # BCa confidence intervals
│   ├── power.py         # Power analysis
│   ├── heterogeneity.py # Stratified IV analysis
│   ├── effects.py       # Effect estimation
│   └── loader.py        # Data loading utilities
└── visualize/
    ├── nature_style.py  # Publication styling
    └── causal_figures.py # Figure generation
```

##### 10.2 Dependencies

- Python ≥ 3.9
- NumPy ≥ 1.21
- SciPy ≥ 1.7
- Matplotlib ≥ 3.5
- (Optional) causal-learn for PC/GES algorithms

##### 10.3 Reproducibility

All simulations use: - Fixed random seed (default: 42) - Deterministic NumPy operations where possible - Version-pinned dependencies

##### 10.4 Computational Requirements

| Study Type | Estimated Runtime | Memory |
|---|---|---|
| Single simulation (200hr) | ~5 min | ~500 MB |
| Baseline validation (20 reps) | ~2 hr | ~1 GB |
| Full parameter sweep | ~8-24 hr | ~2 GB |
| Bootstrap (1000 samples) | ~30 min | ~500 MB |

---


---

*CAUSANTA Supplementary Materials v1.0 For Nature Methods submission*
